# Emergence and robustness of 3D complex tissue architectures from simple physical forces

**DOI:** 10.64898/2026.11.02.763567

**Authors:** Pauline Chassonnery, Jenny Paupert, Anne Lorsignol, Childérick Séverac, Marielle Ousset, Louis Casteilla, Diane Peurichard

## Abstract

Although at central stake in biology and biological engineering, our understanding of how biological tissues acquire their architectures during morphogenesis and which mechanisms have been selected through evolution remain limited. Using adipose tissue as a biological model, we build a 3D mathematical model with simple mechanical rules between cells and fibers, and show that the model reproduces, in-silico, not only the emergence of complex and heterogeneous 3D architectures, but also the intra and inter-individual heterogeneities of shapes as observed in vivo. The model reveals that specific rules for cell differentiation are not necessary for tissue structuring, but could have been selected through evolution for robustness and efficiency. This study therefore suggests a paradigm shift where tissue architecture is an emerging process surprisingly more driven by mechanical processes than by complex predetermined molecular or genetic programs.

**Significance Statement:** Understanding how tissue structure emerges may enable the identification of strategies to generate, preserve or restore its architecture. A simple 3D mathematical model calibrated on biological samples reveals that complex biological tissue organizations can emerge from simple stochastic physical rules, independently of a classical genetic determinism or prior biological knowledge about the final architecture. This view opens new alternatives and perspectives in biological engineering of organoids and organ-on-chip. This also paves the way for innovative physics-based therapies in addition to genetic interventions, and for the development of digital twins.

## Introduction

The overall three-dimensional architecture of an organ is essential to its physiology and health. For all organs or tissues, it generally represents the most efficient configuration for performing specific functions (1,2,3) and any structural alteration inevitably leads to functional decline and diseases (4-9). Understanding how and why these architectures develop has always been a fundamental challenge in biology, medicine and biological technologies (10). For decades, these questions have been explored via reductionist approaches, consisting in breaking down large systems -such as organs-into pieces -such as their individual molecules- and looking for causal relations between the system behavior and its individual parts. These approaches, amplified by the development of increasingly powerful technologies such as omics, have led to a massive accumulation of knowledge at the molecular level and have played an important role in identifying molecular targets that can participate to control proper architecture development (11). However, they also reveal that many factors can operate within redundant and compensatory networks, forged and selected progressively by evolution, leading to an overall complexity which remains difficult to understand/summarize (9).

Therefore, understanding biological systems as a whole requires to modify our current approaches to account for more complex interactions, including feedback loops leading to dynamic features with high-level properties that cannot be deduced from the properties of their individual components alone. This latter vision, called ‘emergence’ and associated with complex systems theory, has recently acquired the status of a ‘new concept’ for biology (12-16). Such a transformation of conceptual approaches has previously occurred in ecology, which underwent a major shift with the adoption of the ‘emergence’ viewpoint, leading to significant advances and a more integrated understanding of ecosystems (17). The traditional linear predator-prey chain has been replaced by a dynamic network of nonlinear interactions in which species influence each other in multiple ways and stability emerges from the interactions of the entire system over time. This paradigm shift has been proposed in other fields such as tissue biology (18) or aging (19) showing the explanatory power of the ‘emergence’ viewpoint compared to the traditional linear approach.

Uncovering the initial rules of tissue architecture emergence could represent a major shift for biology and biological engineering, by the identification of the primary drivers of tissue development. However, this is difficult to assess through experimental observations alone, since the studied systems contain numerous redundant intricated mechanisms shaped by evolution (20). This is where mathematical models can represent an unvaluable tools, by offering an abstract and simplified framework with complete control on the system components and rules. Among them, Individual based models (IBM) offer great flexibility and allow for the integration of multiple scales of organization, discrete local properties, and stochastic events (21-25). Because of these characteristics, IBMs are particularly well suited for exploring how complex tissue architectures can emerge from simple local interaction rules at the cellular level.

Among the simple rules that could structure the organism, the importance of biomechanics has long been recognized by biologists. Many works have focused on mechano-transduction, i.e. the process by which cells adapt to mechanical inputs through mechano-sensing receptors and translate this information into signaling events to trigger biological responses (26-32). It is now well documented that cells are able to adapt to and shape themselves the mechanical environment they evolve into, through a complex machinery involving ion channels, transcription factors, molecular and genetic programs, that have been put in place throughout evolution. These mechanical rules are often considered fundamental, but they are almost always regarded as a direct consequence of a genetic process rather than as the primary cause (33). However, a recent publication challenged this view (34), by showing that simple physical rules are sufficient to explain the skin 2D fragmentation on the head of crocodiles, independently of genetic processes. In this paper, using a systemic approach, we build a 3D mathematical model confronted to experimental data to explore whether complex tissue architectures can emerge from simple mechanical rules between a reduced set of agents, without complex predetermined genetic programs.

In multicellular organisms, many secretory organs – including endocrine organs-have glove-like shapes with specialized cells grouped in clusters around a vascular tree. This is the case in adipose tissue (AT), the largest endocrine organ in mammals. Vital for life due to its role in energy homeostasis, AT is also primarily involved in the global obesity pandemic, a major public health issue (35). Because alterations in its architecture are tightly linked to changes in its mechanical properties (36) and correlate with major metabolic disorders, AT provides a physiologically and clinically relevant model to dissect how cellular and extracellular mechanical interactions drive the emergence of tissue architecture. While the overall glove-like structure of AT is observed for all subunits in all animals, its precise shape exhibits high intra- and inter-individual heterogeneity (37). The overall architecture, very efficient for exchanges, is generally considered as mainly governed by the vascular system. This is supported by developmental studies revealing that adipocyte differentiation is firstly observed as a central cluster located near blood vessels followed by a progressive centrifugal differentiation (38), and consistent with the nutrients supply from the blood stream that play a key role in the differentiation of adipocytes from stromal cells. Moreover, several studies (39,40) showed the location of adipose progenitors close to the vasculature, and explained the adipocytes centrifugal differentiation by the paracrine secretion of adipogenic inducers by differentiating cells (41,42). All these observations led to the commonly accepted paradigm that adipose tissue architecture is primarily guided by the structure of its vascular system. In this paper, we aim to challenge the present understanding by hypothesizing that biological architectures can emerge from simple mechanical rules between few components of the tissue, to which additional mechanisms have been added throughout evolution for robustness and energy efficiency. Using a calibrated 3D computational model and 3D tissue imaging and segmentation, we show that the complex architecture of AT and its associated heterogeneity may not be primarily driven by vasculature, but can emerge from simple mechanical interactions between cells and an extra-cellular matrix, independently of the initial location of these agents. We propose that all events associated with the vascular system were subsequently selected by evolution because they bring robustness and efficiency to the process.

## Results

### 1) *In vivo* inter and intra-heterogeneity of 3D adipocytes clusters

We first imaged large volumes of 8 adipose half pads in healthy adult mice with light sheet microscopy, allowing high-resolution images. To process and analyze these vast amounts of complex imaging data, we developed 3D image processing tools for automatic detection of cell clusters (Fig 1.a and Supp Fig 1) and computed morphological quantifiers such as cluster elongation Elob and solidity Slob (Supp Methods). This analysis revealed between 7 to 21 lobules for each adipose pad for a total of 108 segmented cell clusters. Based on the Elob and Slob quantifiers (Fig 1.b), we classified cell clusters into three main types: Solid Elongated (SE, well packed and elongated structures with large values of Elob and Slob), Solid Rounded (SR, well packed and round structures with low value of Elob and large value of Slob) and Branching clusters (Br, tortuous geometries with low values of Slob). The SE/SR/Br proportion revealed an intrinsic level of heterogeneity in each sample (i.e. intra-individual heterogeneity, Fig 1.a). The comparison of these proportions between samples also revealed an inter-individual heterogeneity (Fig 1.c). All samples combined, we found a large majority of SE lobules, some SR lobules and a few Br lobules (around 60, 34 and 6% respectively, Fig 1.c). Altogether, these data revealed the complexity and the intra- and inter-individual heterogeneity of in vivo cell cluster shapes.

**Figure 1.**
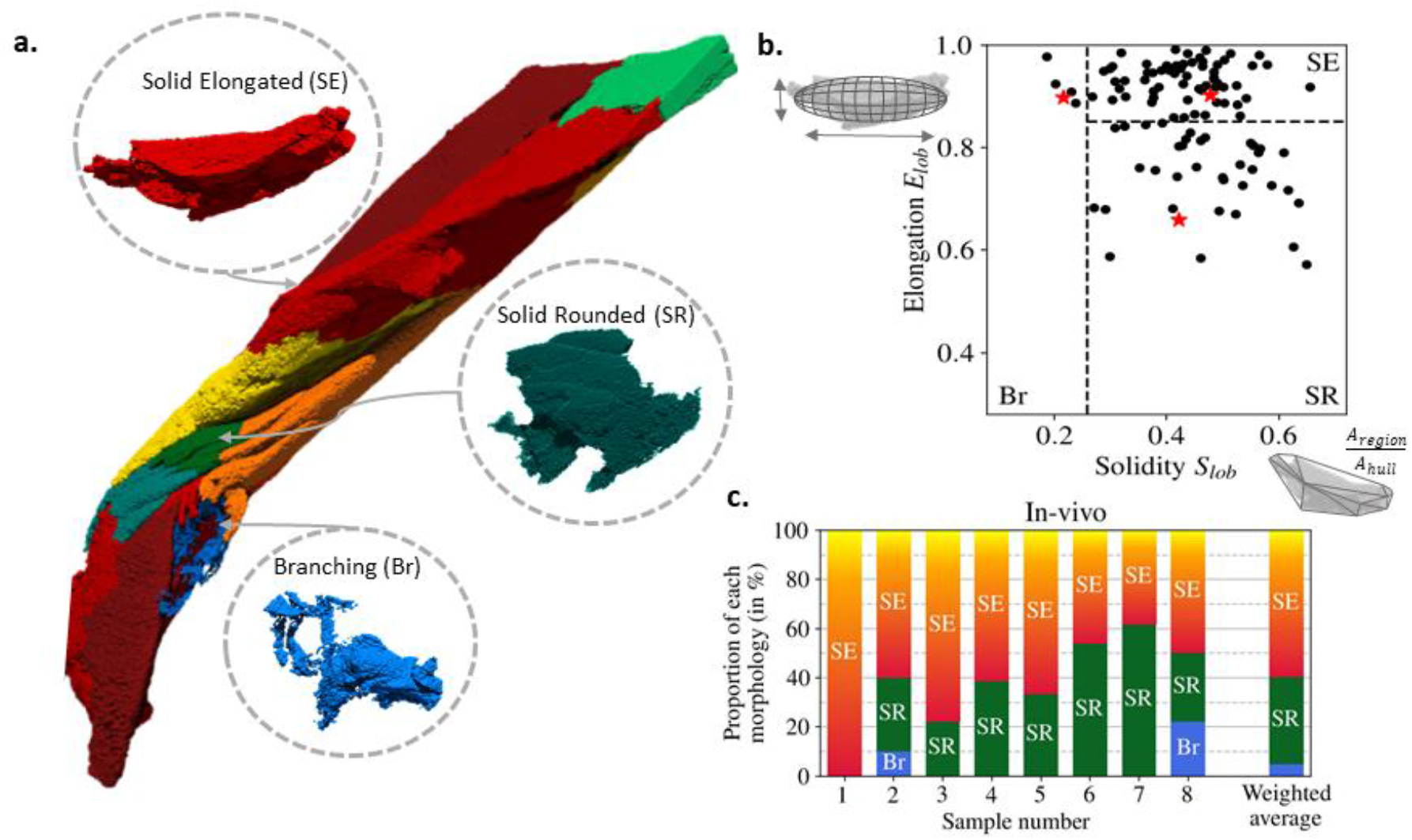
In vivo inter and intra-heterogeneity of 3D cell clusters. **a.** Example of a segmented AT tissue colored by cell cluster types (SE red/yellow, SR green, Br blue). **b**. Distribution of the in vivo cell clusters (total 108 clusters from 8 different samples) according to the morphological quantifiers Elob (elongation, in ordinate) and Slob (solidity, in abscissa). The red stars relate to the three cell clusters highlighted in panel a. **c**. Distribution of SE/SR/Br lobules in vivo for the 8 segmented samples. The weighted average proportion all 8 samples combined is given on the right

### 1) Emergence of realistic tissue architectures using simple mechanical interactions

In order to test the emergence of complex 3D tissue architectures from simple mechanical forces between cells and the Extra-Cellular Matrix (ECM), we constructed a 3D IBM with two agents, adipose cells and an ECM, and as many free parameters as possible. In a cubic periodic domain, the ECM network was modeled as a set of dynamically interconnected cylinders of given length and width (42). These fiber units were initially randomly positioned and oriented in the domain. Along time, fiber units could randomly connect with new intersecting neighbors (probability *X*_*link*_) and existing crosslinks could disappear, to account for the dynamical remodeling of the ECM network. Both processes followed random (Poisson) laws, of frequencies *v*_*link*_ and *v*_*unlink*_ respectively, such that 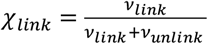 represented the mean proportion of crosslinks among the pairs of intersecting fibers, at equilibrium. Fiber crosslinks were modeled by linear springs forcing the connected fibers to maintain a proximity between their attachment sites during motion, and were supposed to have a constant flexural modulus controlling the resistance to bending of long fibers. Adipocyte cells were modeled as 3D spheres with time-dependent radii. Initially, no cells were present. In time, new cells appeared randomly with a minimal radius *R*_*min*_ and the volume of existing adipocytes grew regularly until a maximal radius *R*_*max*_. Both cell insemination and growth accounted for hyperplasia (differentiation of new adipocytes from progenitors) and hypertrophy (growth of existing adipocytes) respectively.

Cells and fibers repulsed each other to account for competition for space. All mechanical interactions (cell-cell, cell-fiber, fiber-fiber) were modeled by soft potentials to account for small deformations of the elementary units. As a first step, cell insemination occurred randomly in the spatial domain to focus on the role of mechanical interactions on the emergence of complex spatial architectures, independently of a specific rule for cell insemination. All the details of the mathematical model are given in Methods and illustrated in Supp Fig 2, and the model parameters are summarized in Supp Table 1.

We first focused on the role of four free parameters: the volume fraction of fibers *Φ*_*fib*_ (controlled by the total number of fiber), the volume fraction of cells *Φ*_*ad*_ (controlled by the maximal number of inseminated cells), the crosslink proportion *X*_*link*_ and the fiber crosslinking frequency *v link*. Note that the fiber unlinking rate was deduced from the relation 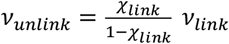. We produced 10 simulations per parameter value to account for stochasticity, for a total of 3240 simulations with these 4 parameters. The simulations were visualized and analyzed effectively using the same segmentation tools we used to analyze the in vivo images. (see Methods and Fig 2.a). The cell clusters were stained and classified into SE, SR, and Br types using the same Elob and Slob morphological quantifiers as those used in vivo (Fig 2.a).

**Figure 2.**
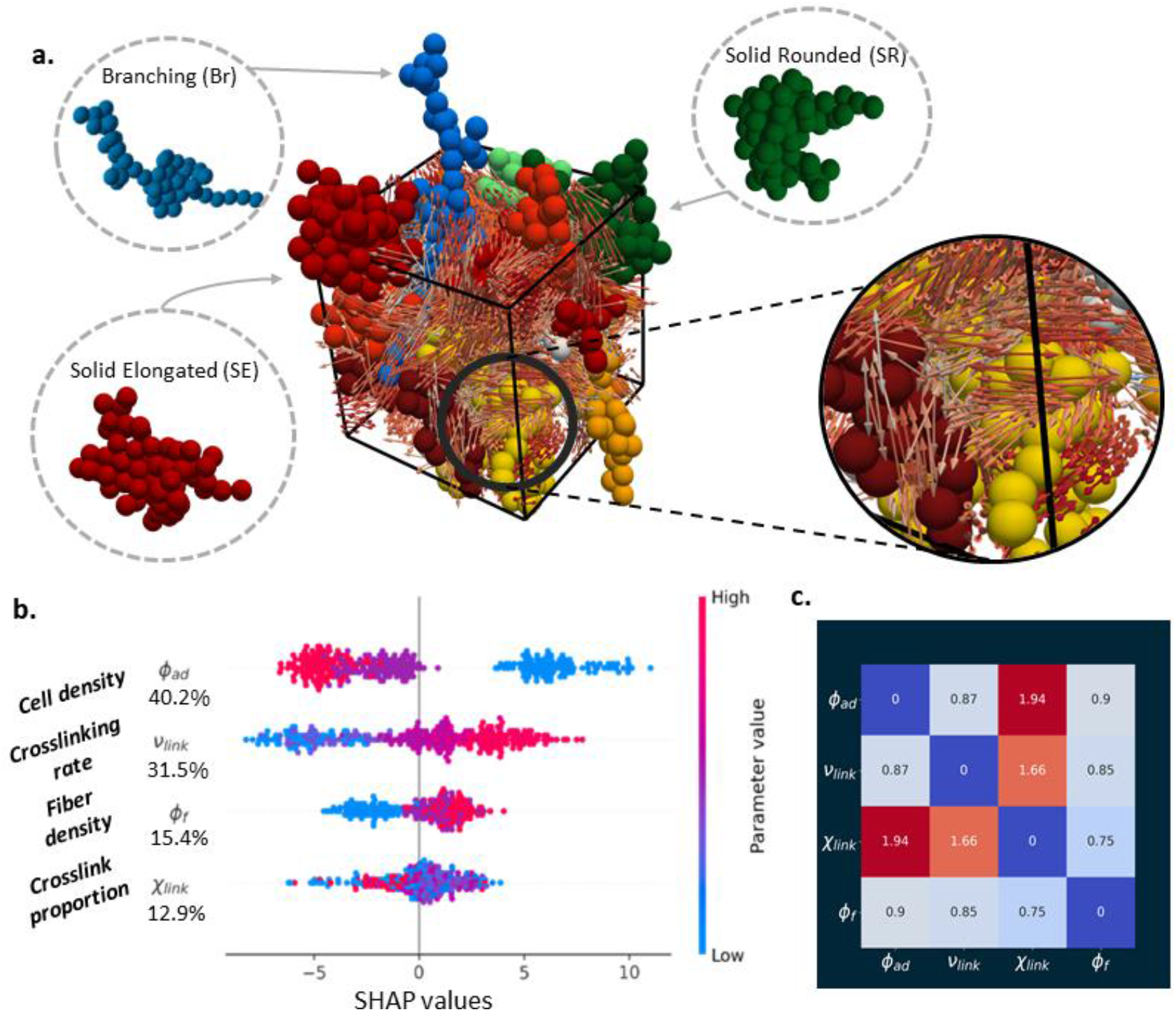
Emergence of realistic tissue architectures with random insemination. **a.** Example of a simulation result at equilibrium in a 3D periodic square domain. Cells are represented as spheres colored according to the cluster they belong to. Fibers (spherocylinders) are represented by their central segment (double arrow), colored according to their local alignment with their neighbors (from blue (low) to red (high) local alignment, visible in the magnification on the right). **b**. SHAP summary plot with parameters ranked by order of importance in the prediction of ΔQsample (distance between In vivo and In silico cluster distributions). The color in each graph represents the value of the associated parameter (red and blue for large and low values of the parameter, respectively). A positive SHAP value means that the parameter increases the prediction of ΔQsample (worsen the in silico – in vivo fit) whereas a negative SHAP value means that the parameter decreases the prediction of ΔQsample (improves the in silico – in vivo fit). **c**. Absolute mean plot highlighting interaction effects (off-diagonal) of the parameters in the prediction of ΔQsample. Colors represent large (dark red) to small (dark blue) effects.

The first surprising result was the ability of the model to generate heterogenous cell clusters, separated by more or less aligned fiber sheets (Fig 2.a). The in-silico fiber structures were efficiently stained and characterized by a quantifier *Al*_*mean*_ measuring the local alignment of the fiber units (see Methods and Fig 2.a). Moreover, the systems at equilibrium showed the co-existence of 3 clusters subtypes revealing an intra-individual heterogeneity similar to the one observed in vivo (Fig 1.a in vivo, Fig 2.a in silico). Among the 73,841 3D cell clusters from 3,240 simulations, 62.75% of clusters correspond to SE-type clusters, followed by SR- and much more rarely Br type clusters (32.67% and 4.58% respectively). Remarkably, these proportions corresponded to those observed in vivo (Fig 1.b in vivo Supp Fig 3.a in silico).

Altogether, these results showed that non-adhesive cells growing in a dynamically reticulated network of fibers can spontaneously lead to the appearance of cell and fiber structures that match in vivo complex architectures, without the need for complex genetic or chemical programs prescribing the differentiation of specific cell types in determined locations of the tissue. It is noteworthy that the *in vivo* lobules had a larger volume than the *in silico* lobules. Indeed, the largest *in silico* lobules had a volume of Vlob = 356 cells ≈ 2.108 *µm*^3^ which was comparable to that of the smallest *in vivo* lobule. Nonetheless, we noted that the difference between the smallest and the largest lobules was of the order of a 1:50 ratio for both the *in vivo* and *in silico* data, indicating that the size variability of our model was comparable to that found in biological tissues.

### 1) Reproducing the in vivo cell cluster distributions requires elevated cell-to-ECM ratio and slow ECM remodeling dynamics

We next investigated whether the model was able to simulate quantitatively well the biological data. To this aim, we calculated an index, Δ*Q*_*sample*_, to measure the distance between the in silico and in vivo SE/SR/Br cluster distributions, taking into account the intra- and inter-individual heterogeneities of the in vivo samples. For each quadruplet of values *P* = (*Φ*_*f*_, *Φ*_*ad*_, *X*_*link*_, *v*_*link*_) tested, we computed:

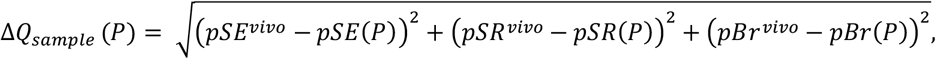

where pSE(P), pSR(P) and pBr(P) represented the average percentage of each cluster types at equilibrium in the 10 simulations for this quadruplet P (ranging from 120 to 250 clusters per parameter combination), and pSEvivo, pSRvivo and pBrvivo represented the percentage of each cluster type computed on the *in vivo* images of all 8 combined samples (108 clusters). Low values of *ΔQsample* indicated proximity in the *in silico* and *in vivo* cluster distributions and therefore good fits, while large values indicated lesser fits. The values of the *in silico – in vivo* distance *ΔQ*_*sample*_ were then represented in the 4D parameter space (*ϕ*_*ad*_, *ϕ*_*fib*_, *v*_*link*_, *X*_*link*_) (Supp Fig 3.b). To gain more quantitative insights into the role of each parameter and their interactions in the *in silico – in vivo* fit, we adopted an explainable Machine Learning (ML) strategy. Simulations were randomly split into training (80%) and test (20%) sets before training a tree-based ML Random Forest (RF) model to predict the values of *ΔQ*_*sample*_ from the parameter values. The model predicted the value of *ΔQ*_*sample*_ with 67% R2-score, 29% NRMS and 3% RRMS errors (see Methods for more details on the scoring measures) on the test set. To define the respective contribution of each variable in the fit predictions, we next applied the Shapley Additive exPlanations framework (SHAP) to our surrogate ML model. The results showed that the cell volume fraction *Φ*_*ad*_ and the ECM crosslinking rate *v*_*link*_ were the most sensitive parameters in predicting *ΔQ*_*sample*_ (with 40.2 and 31.5% importance in the fit prediction respectively, Fig 2.b) while the fiber volume fraction *ϕ*_*fib*_ and ECM crosslinking proportion *X*_*link*_ were less performant (15.4 and 12.9% respectively, Fig 2.b). Moreover, SHAP summary plots (Fig 2.b) revealed that the cell volume fraction *Φ*_*ad*_ was negatively correlated to the *ΔQ*_*sample*_ values, indicating that the best *in silico – in vivo* fits were obtained with a high cell volume fraction. The reverse relationship -to a lesser extent-could be drawn for *Φ*_*fib*_, as more realistic cell cluster distributions were predicted in ECM with low fiber volume fraction. For the ECM dynamics, the better fits with in vivo data (smaller *ΔQ*_*sample*_ predictions) were observed with smaller values of *vlink*, corresponding to ECM with slow fiber crosslinking. Finally, while the ECM connectivity *X*_*link*_ was less important to predict *ΔQ*_*sample*_, this parameter interacted strongly with the main parameters *Φ*_*ad*_ and *v*_*link*_ (Fig 2.c and Supp Figs 3.c-d). Altogether, the sensitivity analysis revealed that the emergence of realistic cell clusters distribution was mainly affected by the number of cells, the ECM dynamics and their interactions with the ECM crosslink proportion.

In conclusion, realistic distributions of cell clusters emerge from the model even when cell insemination is random in space, provided that the system contains a sufficient number of cells and an ECM with a low crosslinking rate.

### 1) In silico intra and inter-individual cluster heterogeneities

We then studied how the two parameters with the greatest impact on *in vivo - in silico* fit (cell volume fraction *Φ*_*ad*_ and crosslinking rate v_link) affected the tissue architecture heterogeneities, for high and low crosslinked fiber networks (*X*_*link*_=0.8, Fig 3 and *X*_*link*_=0.1, Supp Fig 4).

**Figure 3.**
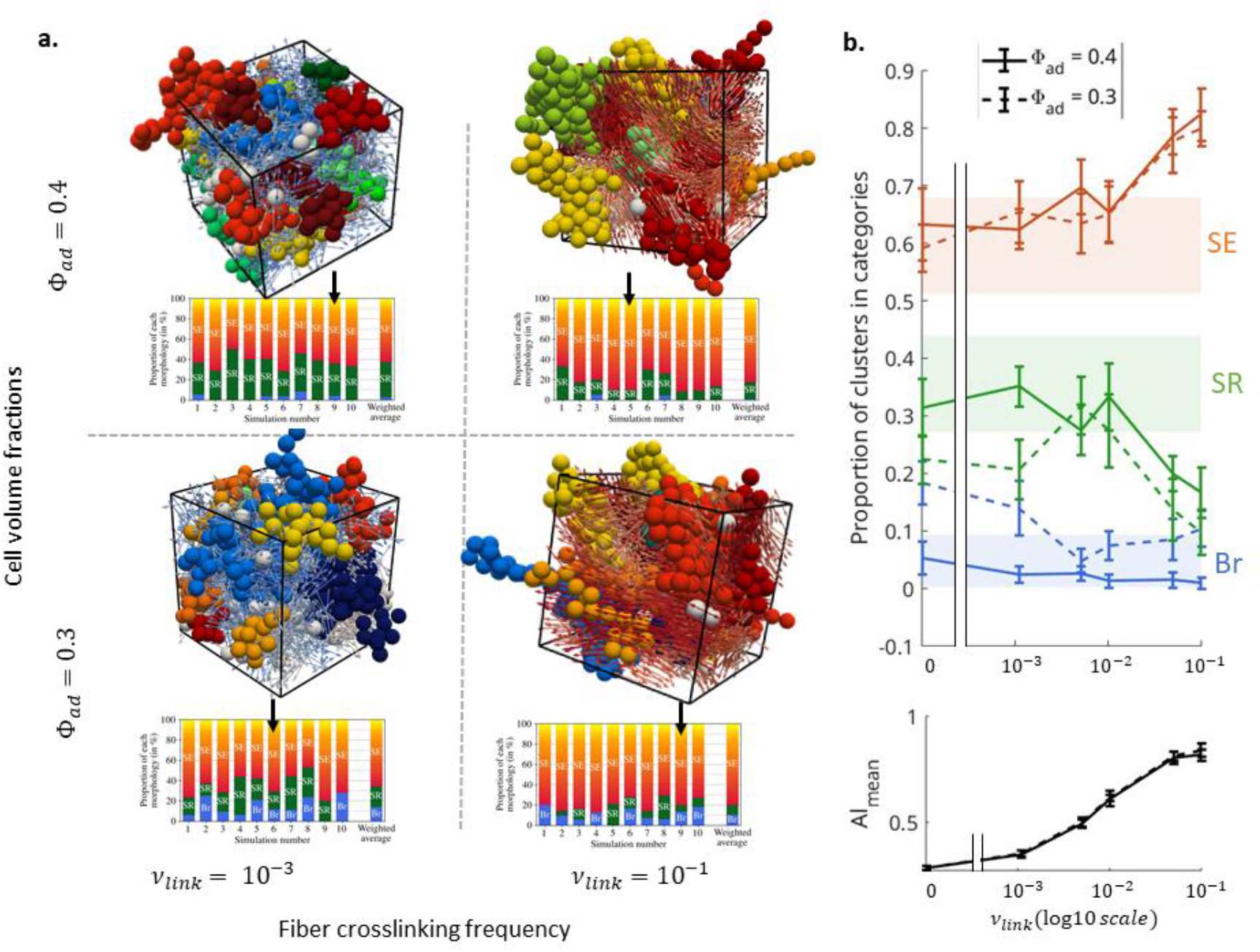
ECM crosslinking drives 3D tissue architecture in silico: sensitivity analysis. **a.** Simulations at equilibrium for *Φ*_*fib*_=0.4, highly crosslinked ECM *X*_*link*_=0.8. top line: *Φ*_*ad*_=0.4, bottom line: *Φ*_*ad*_=0.3. left column: low crosslinking rate *v*_*link*_ = 10^−3^ . right column: fast crosslinking rate *vlink* = 0.1. Cells are colored according to their cluster category (red/yellow for SE, green for SR, blue for Br). Fibers are colored according to their local alignment (from blue (low) to red (large)). The SE/SR/Br proportions for the 10 simulations of the selected parameter sets and their weighted mean are indicated in the bottom bar plots. b. top: mean (weighted) proportions of SE (red), SR (green) and Br (blue) clusters as function of *v*_*link*_ (log10 scale) for *Φ*_*ad*_=0.4 (plain curves) and *Φ*_*ad*_ = 0.3 (dotted curves), and for in-vivo data (shaded error bars). Plain error bars indicate weighted standard deviation between simulations (10 simulations per parameter values). bottom: Fiber mean alignment as function of *v*_*link*_ (log10 scale).

For a high level of crosslinked fiber network ( *X*_*link*_=0.8, Fig 3), we first observed an intrinsic level of heterogeneity in the SE/SR/Br distributions, for a same set of parameters (comparing the proportion bars under the 3D simulations in each panel of Fig 3.a).

Then, comparing the mean SE/SR/Br distributions between parameter sets, we observed higher proportions of SE (elongated) cell clusters in fast crosslinking ECM (*v*_*link*_ = 10^−1^, right panels of Fig 3.a) compared to slow crosslinking networks (*v*_*link*_ = 10^−3^, left panels of Fig 3.a). The highest proportions of SE elongated clusters were also associated with more aligned fibers as revealed by the level of red colored fibers in the two 3D simulations on the right compared to those on the left. Moreover, systems with higher cell volume fractions (*Φ*_*ad*_=0.4, top panels of Fig 3.a) generated fewer Br clusters than systems containing fewer cells (*Φ*_*ad*_=0.3, bottom panels of Fig 3.a). All these observations were quantitatively confirmed by plotting the SE/SR/Br proportions (Fig 3.b, top) and fiber mean alignment Al_mean (Fig 3.b bottom) as function of *v*_*link*_. Indeed, the proportion of SE clusters (red curves in Fig 3.b top) increased with *v*_*link*_, concomitantly with the decrease in SR clusters (green curves in Fig 3.b top). Moreover, a lower cell volume fraction *Φ*_*ad*_=0.3 was associated with a higher proportion of Br clusters (compare the plain and dotted blue curves of Fig 3.b top). Strikingly, the fiber mean alignment *Al*_*mean*_ also increased with *v*_*link*_ (Fig 3.b bottom) independently on the cell volume fraction, showing that, for highly crosslinked networks *X*_*link*_=0.8, fiber alignment was mostly controlled by its crosslinking rate v_link. The intra-individual heterogeneity was also dependent on the ECM characteristics. Indeed, fast crosslinking ECM networks produced systematically more SE clusters, reducing the variability of SE/SR/Br proportions between simulations compared to slow crosslinking networks (compare the error bars for *v*_*link*_=0.1 with *v*_*link*_=0 in Fig 3.b).

We then investigated the case of lowly crosslinked ECM *X*_*link*_=0.1 (Supp Fig 4). In these conditions, the ECM could not reach the high level of alignment observed for *Xlink*=0.8, even for fast crosslinking networks (*Al*_*mean*_ plateaus at ≈0.5 for *X*_*link*_=0.1 and to ≈0.9 for *X*_*link*_=0.8, compare bottom plots of Fig 3.b and Supp Fig 4.b). The lack of ECM alignment ability reflected in the proportion of SE clusters, that remained almost constant as function of *v*_*link*_ for *X*_*link*_=0.1 (bottom panel of Supp Fig 4.b). As previously described for the case with highly crosslinked networks, the proportion of SR and Br clusters depended on the number of cells growing inside the ECM network, with more SR clusters for *Φ*_*ad*_=0.4 compared to *Φ*_*ad*_=0.3.

Altogether, these results showed that the increase of the number of cells in the model induced spontaneously more organized configurations with a clearer segregation between the cells and the ECM network. Cell cluster shapes were strongly influenced by the ability of the subjacent network to deform under an increasing pressure exerted by the growing cells. Less linked ECM or ECM with slower crosslinking were associated with the emergence of architectures with 60-70% SE and 20-30% SR clusters, with high intra-individual heterogeneity. In mirror, highly linked and fast crosslinking ECM produced more elongated and rigid fiber sheets, leading to the emergence of very aligned cell clusters (majority of SE clusters) and less intrinsic heterogeneity.

### 1) Biased insemination is not necessary for cell clusters formation but confers robustness and efficiency

So far, our results showed that cell clusters similar to those observed *in vivo* could appear with random cell insemination in the domain. However *in vivo*, new adipocytes appear in close proximity to existing adipocytes due to the subjacent presence of a vascular system. In order to study the role of a pre-determined structure for nutrient distribution and cell differentiation, we considered a biased insemination process that promoted grouped insemination. To achieve this, the positions of newly inseminated cells were chosen randomly according to a probability law proportional to the local adipocyte density to the power *b*_*ins*_, (supp Fig 2.c). A new set of simulations was performed with *b*_*ins*_ = 0.1 and 0.5, exploring 3 values for *ϕ*_*fib*_ and *ϕ*_*ad*_, 36 pair values of (*X*_*link*_, *v*_*link*_) and 10 simulations per parameter set, giving a large database of 6480 new simulations.

For parameter combinations corresponding to a good fit without bias (low values of *ΔQ*_*sample*_), the bias did not significantly change the in silico in vivo fit (lower right panels of Fig 4.a, corresponding to *ϕ*_*fib*_ < *ϕ*_*ad*_. In contrast, in parameter regions failing to produce realistic cell clusters distribution without bias (upper left panels of Fig 4.a, corresponding to *ϕ*_*fib*_ ≥ *ϕ*_*ad*_), the bias improved significantly the fit (lowered the values of *ΔQ*_*sample*_). To quantify the effect of the bias, we performed a new ML based sensitivity analysis for the new dataset including the bias parameter *bins*. The previous SHAP framework was applied to the new Random Forest (RF) ML model (R2-score 65%, NRMS error 30%, RRMS error 2%) to predict the value of *ΔQ*_*sample*_ on the new parameter set. The summary plot (Fig 4.b) revealed that (*b*_*ins*_, *ϕ*_*ad*_, *v*_*link*_) were the most important parameters for the *in silico in vivo* fit prediction, with similar weights (27.9%, 25,7% and 23.2% respectively), while (*ϕ*_*fib*_, *X*_*link*_) remained secondary as in the case without bias (12,6% and 10.6% respectively, Fig 2.b). We also obtained strong interactions between the parameter *X*_*link*_ and (*ϕ*_*ad*_, *v*_*link*_), as in the case without bias, and observed even stronger interactions between *b*_*ins*_ and *ϕ*_*ad*_ (Supp Fig 5.a). In particular, high values of *b*_*ins*_ tended to improve the fit (lower *ΔQ*_*sample*_) for low cell volume fractions *ϕ*_*ad*_ (Supp Fig 5.b). Moreover, looking at the number of cell clusters (Supp Fig 5.c), we observed that the main effect of biased insemination was to reduce the number of SE and Br clusters, promoting the appearance of more compact cell structuration. Finally, we investigated whether *b*_*ins*_ could affect the time at which the first clusters emerge (Fig 4.c). Indeed, increasing *b*_*ins*_ significantly reduced this time, showing that biased insemination favored an earlier formation of cell clusters. Altogether, these results showed that biased insemination was not necessary for a large range of parameters, but (a) favored the emergence of realistic cell cluster proportions in situations where clustering is made difficult for the cells without bias (low cell volume occupancy) and (b) systematically accelerated the formation of cell clusters compared to random insemination

**Figure 4.**
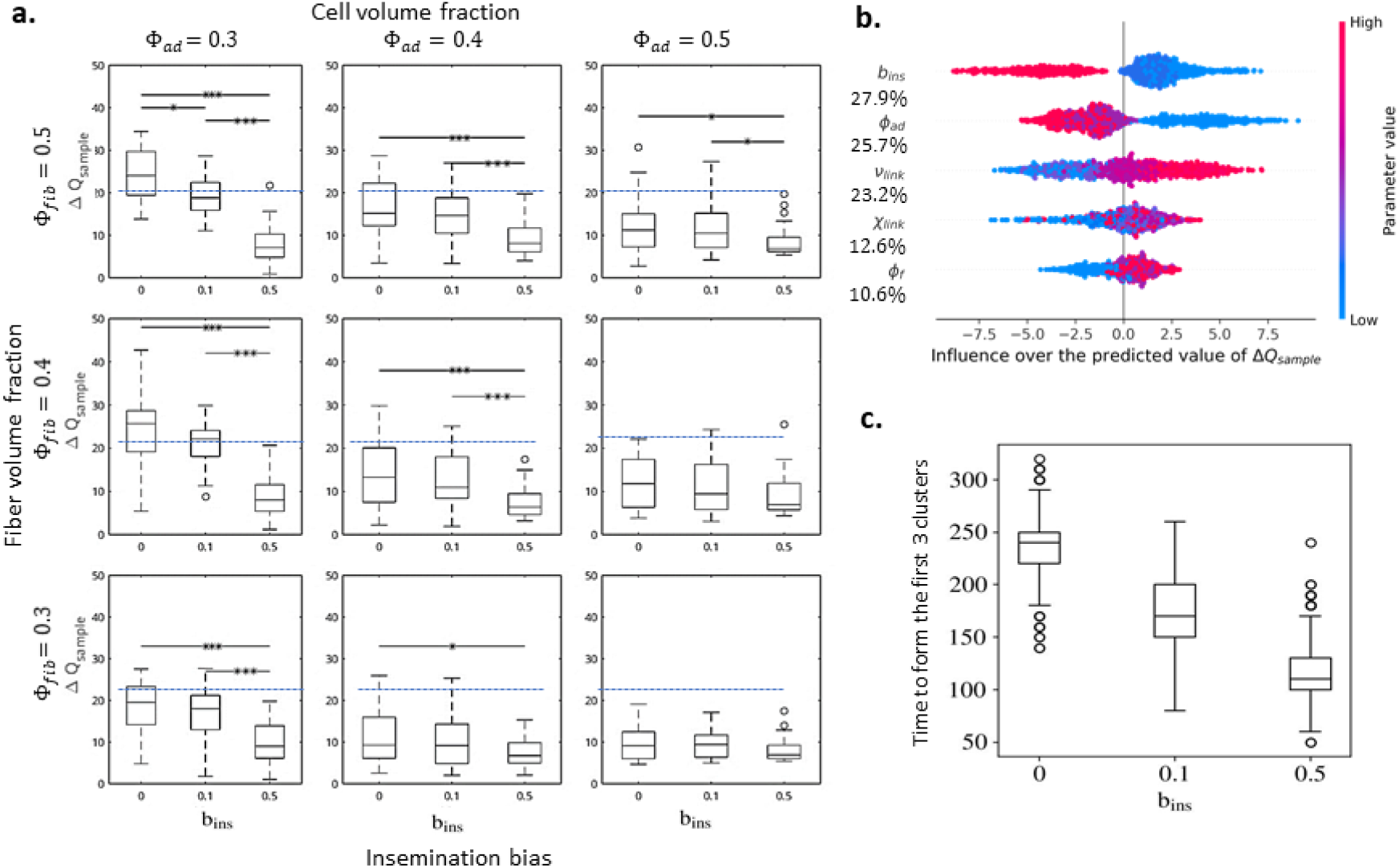
Biased insemination is not necessary for lobule formation but confers robustness and efficiency to tissue structuring. **a.** Boxplots of the in silico – in vivo distance *ΔQ*_*sample*_concatenating parameters (*X*_*link*_, *v*_*link*_) as function of the bias parameter *b*_*ins*_ (abscissa), for different values of *ϕ*_*fib*_ (lines) and different values of *ϕ*_*ad*_ (columns). Statistical differences are measured via Kruskal Wallis tests (***p<0.0001, **p<0.01, *p<0.05). **b**. SHAP summary plot with parameters ranked by order of importance in the prediction of *ΔQ*_*sample*_. The color in each graph represents the value of the associated parameter (red for large values and blue for low values of the parameter). A positive SHAP value means that the parameter increases the prediction of *ΔQ*_*sample*_ (worsen the in silico – in vivo fit) whereas a negative SHAP value means that the parameter decreases the prediction of *ΔQ*_*sample*_ (improves the *in silico – in vivo* fit). Relative importance of the four parameters (*b*_*ins*_, *ϕ*_*ad*_, *v*_*link*_, *X*_*link*_, *ϕ*_*fib*_) in the prediction of *ΔQ*_*sample*_ are indicated as percentages. **c**. Boxplots of the time to form 3 cell clusters as function of the bias parameter *b*_*ins*_ (abscissa), all other parameters combined.

## Discussion

In this study, we propose that short-range mechanical interactions between cells and an ECM fiber network are sufficient to induce the emergence (spontaneous ordering) of a complex 3D tissue architecture, regardless of where new differentiated cells appear. Our model shows that these interactions are sufficient to reproduce realistic tissue architectures, including their intra and inter individual heterogeneities.

This result may seem contradictory to observations made *in vivo*. During the *in vivo* development of the adipose tissue, the complex shape of adipocyte clusters appears to result from the accumulation of newly differentiated adipocytes near existing adipocytes and the vascular tree, suggesting the existence of key local signals for cluster structuring, potentially associated with heterogeneous energy distribution and signals partitioning due to the vasculature (37-41).

In contrast, our data suggest that the structure of the vascular tree and the appearance of new cells near existing cells are not mandatory factors in structuring the tissue. However, these mechanisms -modeled here by a biased insemination process-seem to ensure greater robustness in cluster emergence, by guaranteeing the reproducibility of clusters across a wider range of conditions compared to random insemination. Especially, a biased insemination ensures robustness when cell density is low, situations that occur at the early step of tissue morphogenesis. It also accelerates the appearance of cell clusters, which can be interpreted as greater efficiency. These results suggest that the vascular network structure and the paracrine cell interactions that promote differentiation may not be the primary drivers of tissue development, but could have been selected by evolution for reasons of robustness and efficiency, allowing for a more efficient distribution of chemical energy at specific tissue locations. This new view brings novel perspectives on the function of vasculature and paracrine signaling in tissue morphogenesis.

Our model reveals that the more realistic cell cluster proportions were obtained for small fiber-to-cell volume fractions, lowly connected ECM or highly connected ECM with slow crosslinking. These results are consistent with results in the literature suggesting that real ECM fiber networks do not remodel fast (44), and suggest that the emerging cluster shapes are mainly controlled by the ability of the subjacent network to deform under an increasing pressure exerted by the growing cells. Our model also showed that the emergence of 2D sheets of ECM wrapped around 3D cell clusters does not need a complex geometric representation of the individual fibers. Septa (fibrous sheet-like structures surrounding the lobular cell clusters) simply emerge in our model from mechanical interactions between spherocylindrical crosslinked units (fiber elements) and spherical objects (cells). Another interesting result of the model is that fiber alignment is driven by the parameters related to the connectivity of the network (fiber crosslinking rate and linked fiber fraction) even in the presence of cells. This result is consistent with our previous work on the emergence of 3D fiber network architecture (43), showing that cells have a limited impact on the overall fiber alignment.

Another remarkable feature of our model is that cell-cell adhesion is not required for cell clustering: the spontaneous organization of tissues into cell clusters could be simply the result of competition for space occupancy of growing cells in a mechanically constrained environment. This result is surprising since the evolution of multicellularity is associated with the development of intercellular attachments of individuals cells (45). However, most mammalian cells are covered by the glycocalyx, a carbohydrate-rich layer composed of glycoproteins and glycolipids. Due to its high content of negatively charged residues, such as sialic acids, the glycocalyx generates electrostatic repulsion that prevent non-specific adhesion between cells (46). We suggest that cell adhesion, which is linked to multicellularity, is not essential for tissue structuring but could also be seen as a mechanism conferring robustness and stability to the final structure subject to external mechanical stresses (47).

Remarkably, the model reproduces the intra and inter-individual heterogeneities of the cell cluster distributions as in vivo observed. The fact that different simulations for a same set of parameters produce heterogeneous tissue architectures suggests that at least part of the inter-individual heterogeneity observed in vivo, could simply be due to stochastic variations around the same mechanisms. The in vivo definite demonstration of such conclusion is often impossible but stochastic events including cell specification have often been described and associated with robust outcomes (48). This suggests that a part of the interindividual differences can never be reduced and controlled, a point to take into account in biological engineering and personalized medicine.

Finally, on an energetic viewpoint, it is noteworthy that the more realistic architectures correspond to systems with the lowest chemical energy spent. Indeed, low-dynamical networks do not remodel their links often, therefore we can reasonably consider that the energy spent for renewal of their crosslink configuration is low. Furthermore, networks with low linked-fiber fraction, even dynamical, do not remodel many links so the total energy devoted to fiber remodeling is also low. In addition, biased/localized cell insemination, which speeds up the process, also offers an energy advantage.

Altogether, these viewpoints challenge the classic genetically or chemically oriented viewpoints where tissue morphogenesis is seen as the result of well-tuned genetic differentiation programs, and where mechanics is either disregarded or seen as secondary/regulatory factor. Here, we propose a counterpoint where mechanics is at the heart of tissue architecture emergence, and where chemistry/genetics could be seen as means for systems to stabilize and consolidate the processes, subsequently selected during evolution. This view also open new alternatives and perspectives in biological engineering, to induce and control the emergence of biological objects including organoids and organ-on-chip rather than attempting to reproduce the tissue in its final architecture as observed in vivo. Overall, this work proposes a paradigm shift for the development of biological tissues, where beyond the genetic dogma pattern, formation/tissue architecture emergence is seen as self-organization structuring driven by generic physical rules reminiscent of the rules driving non-living matter (49-54). In addition to genetic interventions, this also paves the way for innovative physics-based therapies and for the development of digital twins.

## Materials and Methods

### Samples and image acquisition tools for in vivo clusters

Samples were obtained from 7-week-old male C57BL/6 mice obtained from Envigo Laboratories. The samples were embedded in 1% LM agarose before being dehydrated in a 100% methanol (MeOH) solution with agitation. The dehydration solution was renewed every 8 hours until there was no more exchange in the medium. After dehydration, and to initiate transparency, the samples were placed in a solution of 1 vol Benzyl Alcohol (BA); 2 vol Benzyl Benzoate (BB); 3 vol MeOH, for 30 minutes at room temperature, with stirring. The solution was replaced with a solution of 1 vol BA; 2 vol BB and renewed every 8 hours until there was no more exchange in the medium.

Transparent samples were imaged using light sheet microscopy (LightSheet Z7 – Carl Zeiss) and Zen Black 3.2 software. They were glued to a sample holder and immersed in an imaging chamber filled with clearing solution. The light sheet was formed by 2 illumination objectives (5x/0.1foc) located on either side of the sample. The sample was illuminated with the follow laser/filter combinations: 488nm (BP505/545) and 561nm (LP585). The fluorescence signal is collected by a 5x/0.16 (EC-Plan Neofluar, nd = 1, WD = 5,1mm, Carl Zeiss) objective, and dual camera (PCO edge 4.2 sCMOS). The illumination and detection objectives are corrected to a refractive index of 1.56. A computer magnification of 1 was applied to give a resolution of 0,92µm/pixel in X and Y dimension and 10µm/pixel in Z dimension. In order to image the entire region of interest, acquisitions were performed using the ‘Tiles – Bounding Grid’ module with a 10% overlap.

### Segmentation tools for in vivo and in silico clusters

The various stages of the analysis pipeline - image preprocessing, adipose tissue segmentation, lobule segmentation and image stitching - were carried out using Fiji (55,56) and Cellpose (57,58) softwares.

### In vivo image processing

Image opening and pre-processing – For each condition and each sample, we used Bio-Format plugin as implemented in Fiji, in order to open 3D images. We first corrected fluorescence images from noise using a 3D-Median filter (1x1x1 ellipsoidal kernel). Then, images were corrected from background using a rolling ball algorithm (59). For each pixel, this function determines local background value by averaging in a large ball (1000 pixel) all values around the pixel, and subtracts this average value from the original image. <u>Adipocyte segmentation</u> – Automatic detection of adipocyte was carried out using cellpose 2.0 software. Because of the poor segmentation performances of the cellpose-implemented models, we choose to fine-tune our own ML-model using the Human-in-the-loop training function (60). From each preprocessed image, adipocytes were first detected using the Cyto2 model implemented in Cellpose. Then, we manually annotated, one by one, adipocyte not detected or only partially detected by the Cyto2 model (False Negative). For each image, we also took care to remove all false detections (False Positive). Thus, correctly annotated, these images were used again in the Cellpose 2.0 software to fine-tune the initial model (Cyto2) and generate our own model. From each sample and each slice, we used our Cyto2 fine-tuned model implemented in the Cellpose Fiji-wrapper and computed a 2d-segmentation map as a mask output of each detected adipocyte. <u>Lobule segmentation</u> - Then, each 2d-segmentation map was converted as a binary map and inverted, in order to keep area corresponding to adopicyte and lobule boundaries (cytoplasmic membrane and spans). Finally, we filtered each 2-binary map using area parameter as a threshold value (> 5000 pixels) in order to keep lobule boundaries only. <u>Image stitching -</u> We used Stitching plugin as implemented in Fiji software (55) in order to reconstruct the whole adipose tissue stack from the individual input stack.

### In silico image processing

To analyze the spatial structuring of the tissue produced by the model, we developed a fully-automatic segmentation tool to class cells into clusters. Cell clusters were defined based on (i) spatial proximity of cells inside one cluster and (ii) the presence of fibers in-between two clusters. A black-and-white image was created from a simulation, where adipocytes appeared in white and fibers in black (possible intersections between a cell and a fiber being also in black). After cleaning and pre-processing the image, we applied a watershed segmentation algorithm (61,62) to divide it into different regions. A unique cluster number was then allocated to cells belonging to the same region, taking into account periodic boundary conditions. We used the same watershed segmentation algorithm to identify cell clusters in tri-dimensional images of biological tissue samples based on the same two criteria. This enabled a comparison between in vivo and in silico clusters. We used the software Paraview (63) to visualize in 3D the states of the segmented systems (both in vivo and in silico), with cells colored according to the cluster their pertain to.

### Morphological quantifiers

We defined quantifiers allowing to quantitatively describe the local and global organization of the agents observed in the system. We first describe how we quantify the local alignment of the fibers by computing the fractional anisotropy of the fiber directional vectors in a neighborhood of fiber k, then present two quantifiers enabling us to characterize the shape of the lobules.

### Local fiber alignment

In healthy adipose tissue, the ECM fibers form long, aligned bundles that surround the lobules. To check if and how much this kind of organization is present in our results, we defined a numerical quantifier for the local alignment of the fibers, which we call the alignment indicator. Let Ralign denote the sensing distance up to which a fiber may interact with its neighbors: in our models, it is equal to L_fib+d_link^max. For any fiber k, we define its interaction neighborhood Bk as the set of all fibers located within a distance Ralign, plus itself, and its local alignment indicator Alk as the fractional anisotropy of the fibers’ directional vectors within Bk.

It is computed as follows. We denote by *p*_*m*_ = *ω*_*m*_ ⊗ *ω*_*m*_ the projection matrix on the directional vector of fiber m. The mean of the projection matrices of the fibers in Bk is thus given by:

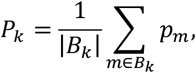

where |Bk| is the number of fibers in Bk. The matrix Pk is symmetric positive-definite, so its three eigenvalues *λ*_1_(*P*_*k*_), *λ*_2_(*P*_*k*_), *λ*_3_(*P*_*k*_), are positive real numbers. The alignment indicator or fractional anisotropy in the neighborhood Bk is then equal to:

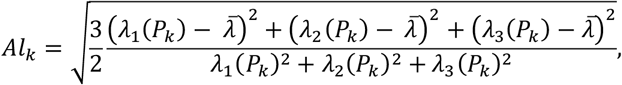

with 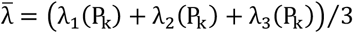 the mean of the eigenvalues. Note that *Al*_*k*_ = 1 if all the fibers in *B*_*k*_ have the same directional vector. If the directional vectors are uniformly distributed then theoretically *Al*_*k*_ = 0, but in practice this is not always the case: indeed, the actual sampling of a random distribution may not be fully isotropic, especially if the number of elements in the sample is small.

To assess the alignment state of the fiber network in our different simulations, we computed the mean of the local alignment indicator Alk over all the fibers of the system and, to account for stochastic variability (due to the random initial condition and the various stochastic processes), we averaged it over 10 simulations conducted with the same set of parameters. We denoted this mean local alignment by Almean.

### Lobule elongation and solidity

To enable a systematic classification and comparison of the lobules segmented from both in vivo and in silico data, we wanted to characterize to their size and shape as simply as possible. To that end, we did not take into account the possible digitations and focused on the core of each lobule. Moreover, though in vivo lobules are polylobed, it has been observed that all the lobes of one lobule tend to be arranged in the same direction. The general shape of its core is therefore a good first approximation for describing the shape of the whole lobule.

A criterion commonly used by biologists is the elongation of the lobules, which can be computed in several manners. Considering that our in-silico lobules were composed of a relatively small number of cells, most of these manners would have been very sensitive to the presence of peripheral digitations or surface irregularities: this was not what we were interested in. In consequence, we chose to fit each lobule with an ellipsoid and computed the elongation (or eccentricity) Elob of that ellipsoid. This was done using Matlab ‘regionprops3’ function, which performs an ellipsoidal fit on each region of the segmented lobule map (see description of the segmentation tools above) and compute many characteristics of the regions and ellipsoids such as their volume, center-of-mass, orientation, elongation, etc.

One of these characteristics is the region solidity Slob, which measures its deviation from convexity. It is expressed as the ratio between the volume of the region and the volume of its convex hull (i.e. the smallest convex polygon containing the region). This quantifier is useful to distinguish between lobules with similar elongation: it especially allowed us to identify, in our in-silico data, lobules with loose, non-compact and convoluted shapes (Supp Fig 2.a).

We thus characterized our lobules using two scalar quantities: their elongation Elob∈[0,1] (equal to 0 if the lobule as no preferred direction and to 1 if it is strictly unidirectional) and their solidity Slob∈[0,1] (equal to 1 is the lobule is convex).

We observed on Supp Fig 2.b that the distribution of the lobules generated by our numerical simulations in the plane Slob – Elob displayed a single, very large peak centered on Slob = 0.34 and Elob = 0.9, meaning that the vast majority of our *in silico* lobules are moderately solid and very elongated. Based on this repartition, we chose to classify the lobules into three categories: (i) branching lobules (denoted as “Br”) for Slob < Scritic = 0.257, (ii) rounded solid lobules (denoted as “SR”) for Slob ≥ Scritic and Elob < Ecritic = 0.851, and (iii) elongated solid lobules (denoted as “SE”) corresponding to Slob ≥ Scritic and Elob ≥ Ecritic. The limits between these three categories are drawn on Supp Fig 2.b in black dashed lines.

### Machine learning analysis

For the random insemination case, the dataset was composed of 3240 simulations from 324 combinations of parameters and for the biased insemination case the dataset was composed of 9720 simulations from 972 combinations of parameters. For each, the datasets were split into train (80%) and test (20%) sets. A Random Forest Regressor (RF) was used and compared to Gradient Boost and Elastic Net regressors, to predict the value of the in silico – in vivo distance *ΔQ*_*sample*_. To better explain the prediction results, a SHAP model was applied to calculate the influences and interactions of each parameter on the output of the predictive model. RF analysis were performed using Scikit-learn, seaborn and shap libraries, and the following Random Forest regressor hyperparameters were used: *n*_*splits*_ = 5, *n*_*estimators*_ = 100, criterion=‘squared_error’. Model performance was assessed using R2-score, Normalized Mean Square Error (NRMSE), and Root Mean Square Deviation (RRMSD).

### Statistical analyses

The study was conducted on Adipose Tissue samples from 8 adult mice. Measurements were taken from distinct samples. All results are given as weighted means ± unbiased weighted SD for the curve plots of cluster proportions (weighted by the number of clusters per sample, Fig 3.b and Supp Fig 4.b), and median ± min/max for the boxplots (*ΔQ*_*sample*_ for Fig 4 and number of clusters for Supp Fig 5.c). Statistical differences were measured using Kruskal-Wallis tests (two-sided) to determine statistical differences between each group (Fig 4.a). The following symbols for statistical significance were used throughout the manuscript: ***p<0.0001, **p<0.01, *p<0.05

### Model

The 3D model featured insemination and growth of spherical cells in a network of dynamically interconnected fibers, based on our previous model of ECM^2^. Given a spatial domain Ω ⊂ ℝ^3^ with periodic boundary conditions, the N_fib_ fiber elements were represented by spherocylinders of uniform fixed length L_fib_ and radius R_fib_, centers **X**_k_(t) ∈ Ω and non-oriented directional unit vectors 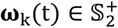. The N_ad_(t) adipocyte elements present in the system at time t are represented by spheres of centers **Y**_i_(t) ∈ Ω and radius 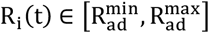. All elements repel their close neighbors by means of a soft repulsion mechanism modeling steric repulsion. Fiber elements have the ability to link to or unlink from each other to model fiber elongation or rupture. The linking and unlinking of fibers follow random (Poisson) processes in time. Fibers also offer resistance to bending through an alignment torque acting between two linked fiber elements. New adipocytes of minimal radius are added to the system following a random process in both time and space, until the maximal number of adipocytes is reached. Existing adipocytes undergo a linear volume growth at constant speed until they reach their maximum radius. Fiber and adipocyte motion and rotation is given by Newton’s second law of motion, in an overdamped regime to model a medium with low Reynolds number.

#### Computation of the mechanical forces

Repulsion: The adipocytes modeled here are physical objects occupying a given volume from which other objects are excluded. Adipocytes undoubtedly classify as deformable objects because their cell walls, cytoplasm and triglyceride droplets are elastic (with an average overall cell stiffness lesser than 1 kPa *(35)*), so that two adipocytes coming into contact will deform and flatten against each other. We can thus use Hertz’s theory for non-adhesive elastic contact between two spheres *(25, 26)* to model the pairwise interaction force 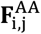 exerted by an adipocyte j on its neighbor i:

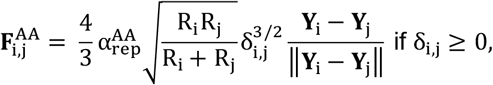

where 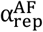 is the maximal intensity of the adipocyte-adipocyte repulsion and *δ*_i,j_ = R_i_ + R_j_ - ||**Y**_i_ – **Y**_j_|| is the depth of the overlap between the two adipocytes.

Just as adipocytes, the fibers in our model are treated as deformable (although rather stiffer) objects. To model the contact between a (spherical) adipocyte and a (spherocylindrical) fiber, we assumed that its geometry was similar to that of a contact between two spheres. This is obviously exact if the contact happens at one of the hemispherical ends of the fiber, and is a reasonable approximation if the contact happens along the cylindrical body of the fiber. Therefore, fiber k exerts on adipocyte i a repulsive force equal to:

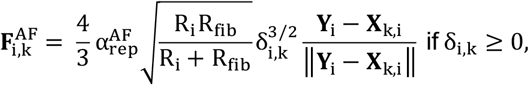

where 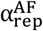 is the maximal intensity of the fiber-adipocyte repulsion, □_i,k_ = R_i_ + R_fib_ - ||**Y**_i_ – **X**_k,i_|| is the depth of the overlap between the two agents and **X**_k,i_ is the point on the central segment of fiber k that is closest to the center of adipocyte i, that is the orthogonal projection of **Y**_i_ onto the segment [**X**_k_ − **ω**_k_L_fib_/2, **X**_k_ + **ω**_k_L_fib_/2] :

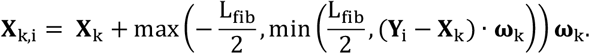

Conversely, adipocyte i exerts on fiber k a repulsive force equal to 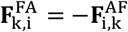 according to Newton’s third law. This force is applied at point **X**_k,i_, thus inducing a rotational torque with respect to the fiber’s center-of-mass:

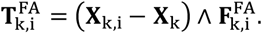

Lastly, the interaction force between two fibers k and m is approximated by the force between two perpendicularly crossing cylinders of radius R_fib_:

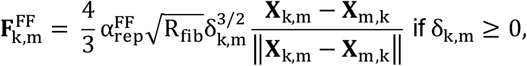

where 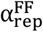 is the maximal intensity of the fiber-fiber repulsion, *δ*_k,m_ = 2R_fib_ - ||**X**_k,m_ – **X**_m,k_|| is the depth of the overlap between the two fibers and **X**_k,m_ and **X**_m,k_ are the closest points along the central segment of both fibers (see appendix A.3 of *(24)* for the actual computation of these points). This force is applied at point **X**_k,m_ and induces on fiber k a rotational torque with respect to its center-of-mass:

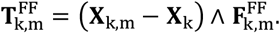

Crosslink forces: Fibers closer than a given threshold 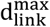 can create a crosslink, modeled as a linear spring of stiffness α_rest_ and unloaded length 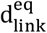 fixed to the two points of the crosslinked fibers that were closest at the time of its creation. Using Hooke’s law, the elastic restoring force sustained by fiber k due to its link with fiber m reads:

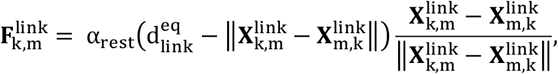

where 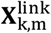 denotes the point of fiber k’s central segment that was closest to fiber m at the time of the link creation. This force induces a rotational torque on fiber k:

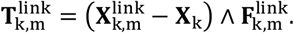

To ensure coherence between the different features of the model, we require that 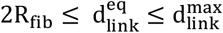.

Furthermore, the links are characterized by a stiffness parameter *α*_align_ > 0 playing the role of a flexural modulus: the larger *α*_align_, the more rigid the fiber network. Given two linked fibers k and m, the additional torque sustained by the fiber k is such that, ∀**u** ∈ ℝ^3^,

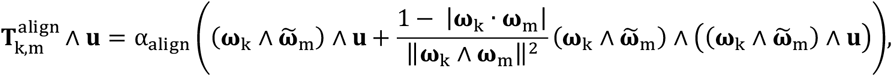

where 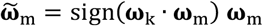 so that there is no preferential orientation.

Friction: We assume that the friction sustained by an adipocyte cell (respectively a fiber) follows a Stokes Law with friction coefficient *µ*_ad_ (respectively *µ*_fib_) *(36)*. The total friction force sustained by an adipocyte i, computed by integrating this law on the whole radius of the cell, is equal to:

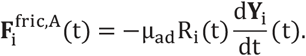

Similarly, the total friction force sustained by a fiber k is equal to:

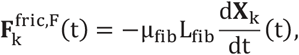

and the associated rotational torque is equal to:

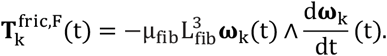

#### Equations of motion

The motion and rotation of all agents can be deduced from Newton’s equation of motion in an over-damped regime. We denote by (p_k,m_(t))_k,m_ the fiber connectivity matrix, that is p_k,m_(t) is equal to 1 if fibers k and m are linked at time t and to 0 otherwise. The motion and rotation of a fiber k ∈ ⟦1, N_fib_⟧ are then given by:

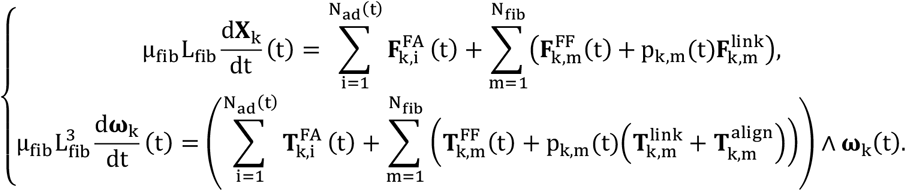

Likewise, the motion of an adipocyte i is given by:

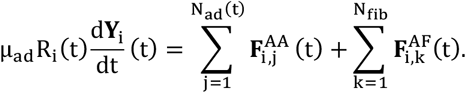

#### Biological phenomena

The main role of AT is to maintain the body energy balance by storing excess energy (carbohydrates and lipids) as triglyceride droplets inside adipocyte cells and releasing it according to metabolic needs. Since we wanted to model the emergence of AT architecture, we were specifically interested in AT morphogenesis, that is the creation of a new layer of adipose tissue to store a continued (not occasional) excessive influx of energy. For this reason, we only included in our model energy storage and not energy release. This storage has two direct consequences: adipocytes increase in volume as they store more fat (hypertrophy), and pre-adipocyte cells differentiate into full-fledged adipocytes (hyperplasia).

Adipocytes differentiation: We account for the latter phenomenon by inseminating new adipocytes with a small radius R^min^ according to a spatial Poisson process with frequency 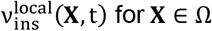, such that 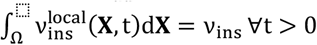. This frequency depends on the local density of existing adipocytes *ρ*_ad_(**X**,t), computed using the particle-in-cell method:

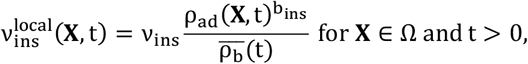

where *v*_ins_ is the global frequency of insemination, b_ins_ > 0 a power factor indicating the magnitude of the dependency and 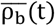 a normalization factor equal to the average of 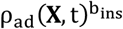 over Ω. Note that, if b_ins_ = 0, all positions are equiprobable, which is equivalent to assuming a uniform repartition of the progenitor cells. If b_ins_ > 0 on the other hand, the probability to inseminate a new adipocyte is higher near existing ones, implying that the progenitor cells are clustered together, mimicking the existence of a vascular network.

Adipocyte growth: We model adipocyte growth, in a simplified way, by a linear volume growth of constant rate Kgrowth capped by the maximal adipocyte radius 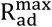:

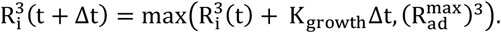

To avoid overcrowding the modeled tissue, we set a maximal adipocyte density *ϕ*_ad_ (equivalent to a maximal domain occupancy rate) from which we determine the maximal number of adipocytes allowed in the system:

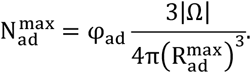

We stop the insemination process when N_ad_(t) reaches 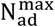.

#### Calibration of the model parameters

The physical scaling of all the parameters of the model, as well as the values used in the simulations, are described in Extended data Table 1. These values are given using a reference length scale U_l_ and time scale U_t_, whose equivalence in physical units are discussed below. A few biologically relevant points may be noted.

Agents’ sizes: The average diameter of a fully grown white adipocyte in the subcutaneous depot is 100 *µ*m. We used in our simulations 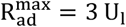, meaning that the reference length scale can be estimated as U_l_ = 33 *µ*m. We set the width of the fiber units to 2R_f ib_ = 1 U_l_ = 33 *µ*m, in agreement with the measures of the width of collagen fibril bundles in loose connective tissues such as adipose tissue *(43)*.

In our model, the flexibility of the fiber network is entirely due to the crosslinks since the fiber units themselves are treated as rigid. Thus, in order for our network to be able to bend around the adipocytes like actual collagen fibers, the length of the fiber units should be about the diameter of an adipocyte. However, achieving the required fiber density with such short fibers would imply to use a very high number of agents, leading to intractable computational costs. We therefore compromised by setting the length of the fiber units to L_fib_ = 6 U_l_ = 200 *µ*m, that was twice the size of fully grown adipocytes.

As in *(24)*, we set the crosslink equilibrium length to 2R_f ib_ so that the link tried to keep the bodies of the spherocylinders touching. On the contrary, we extended the perception distance for link creation to a value slightly larger than its minimum acceptable value of 2R_f ib_. This allowed fibers to link together even if the pressure exerted by the adipocytes tended to move them apart. In this regime, the presence of links therefore participated in a bundling but non overlapping configuration of the fibers.

Agents’ densities: We denote by □_ad_ the maximum adipocyte density of the modeled system, that is the ratio between the maximum volume of adipocytes (after full insemination and growth, without overlapping) and the volume of the spatial domain Ω. Similarly, we denote by *ϕ*_fib_ the fiber density of the system. For each of these densities, we tested three values: 0.3, 0.4 and 0.5. This means that the total agent density *ϕ*_tot_ = *ϕ*_ad_ + *ϕ*_fib_ varied from 0.6 to 1.

These quantities can be compared to the optimal packing density, that is the maximal fraction of the domain that can be occupied by densely packed, non-overlapping agents. For spheres of uniform radius, this optimal packing density is 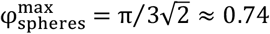. For identical spherocylinders of aspect ratio r = L_fib_/(2R_fib_) = 6, it is equal to:

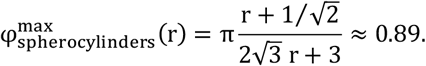

This means that the systems we studied ranged (after full insemination and growth) from rather dense to positively overcrowded, guaranteeing a high level of mechanical stress.

Adipocytes insemination and growth: We set the insemination frequency to 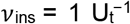 and the adipocyte volume growth rate to 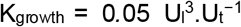. This means that a new adipocyte is inseminated every 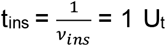 on average, and subsequently takes 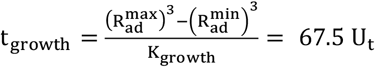 to grow up to its maximum growth radius. These numbers were difficult to relate to actual biological time scales because adipocyte growth rate and differentiation rate are highly variable from one individual to another, depending on many factors such as age, sex, diet and body mass index but also on an innate tendency to favor either adipocyte hypertrophy (growth) or hyperplasia (insemination). However, it has been shown that adipocyte turnover (i.e. the proportion of the total number of adipocytes renewed each year) was relatively homogeneous between individuals independently of all the above-mentioned factors, and was approximately equal to 10% per year *(39, 40)*. Given that the total number of adipocytes in our simulations varied between 573 (*v*_ad_ = 0.3) and 955 (*v*_ad_ = 0.5), a turnover of 10% per year would be roughly equivalent to the production of 100 adipocytes per year, that is t_ins_ = 1 U_t_ ≈ 3 days. This value will serve as a reference time scale for the present study.

Fiber network remodeling: The fiber network could be regarded as having undergone significant remodeling when 10% of its linkable configurations (i.e. existing links and non-linked neighboring fiber pairs) have been renewed through link creation and/or destruction. Given that each configuration is remodeling independently of the others, and considering that *v*_link_ and *v*_unlink_ are of the same order of magnitude, a tenth of the crosslinking processes activated in t_remodeling_ = 0.1/*v*_link_ on average. Excluding the extreme case of non-dynamical networks (*v*_link_ = 0), in our simulations this remodeling time ranged from 3 days (*v*_link_ = 0.1 U_t_^−1^) to 300 days (*v*_link_ = 0.001 U_t_^−1^), which is around the estimated ECM remodeling time of 15 days *(41)*.

Friction coefficients: For the sake of simplicity, we assumed a single friction coefficient *µ*_fib_ for the non-spherical fiber elements of our model. However, we note that several methods for computing the hydrodynamic properties of rigid macromolecules have been developed in the literature. In *(42)* for instance, the authors present a systematic method for computing the frictional tensor of rigid particles modeled as assemblies of beads.

In the absence of empirical data concerning the dynamic viscosity of adipocytes and ECM fibers, we make the simplifying assumption that *µ*_fib_ = *µ*_ad_ = 1 U_m_U_l_^−1^U_t_^−1^.

Total time of simulation: Finally, the total time of simulation T_final_ = 2000 U_t_ was chosen to be at least twice as long as the total adipocyte insemination time 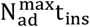. In contrast, it was too short for the fiber networks, except the fastest remodeling ones, to fully equilibrate. This choice was made because long time-scales (compared to the total adipocyte insemination time) were more related to tissue homeostasis than tissue morphogenesis, which was the subject of the present study.

Size of the domain/lobules: As shown in Results, the model systematically creates smaller cell clusters than *in vivo*. We did not expect to be able to compare the volume of the *in silico* and *in vivo* clusters given that the spatial domain used for the simulations has a side length equivalent to 1mm whereas the biological samples are approximately 8 times larger. We note that the small size of the computational domain is a limitation of this study but a necessary choice considering the computational cost of our model. An important validation step would be to reproduce these results in a larger spatial domain and to compare them to more numerous biological samples.

## Data and materials availability

The computer code (Fortran90) used to perform the simulations and the data generated from these computations can be found at https://github.com/chassonnery/3D_AdiposeTissue. The segmentation scripts (in Python) and the Paraview macros automatizing the plotting processes can be made available upon request to the second corresponding author. Source files for the adipose tissue images can be made available upon request to the first corresponding author. All other data needed to evaluate the conclusions of the paper are available in the main text or the supplementary material.

## Acknowledgments

This project was supported by Agence Nationale de la Recherche (ANR) under the project grant number ANR-22-CE45-0024-01. The authors warmly thank Julie Batut, Etienne Danchin and Pierre Degond for fruitful discussions, their careful reading of the manuscript and their excellent suggestions.

## Notes

### Competing Interest Statement

The authors have declared no competing interest.

## References

1. R. N. Mitchell, F. J. Schoen, “Functional Tissue Architecture, Homeostasis, and Responses to Injury” in Biomaterials Science, W.R. Wagner, S.E. Sakiyama-Elbert, G. Zhang, M.J. Yaszemski, Eds. (Academic Press, ed. 4, 2020), pp. 683–700.

2. J. Zhong, W. Huang, H. Zhou, Multifunctionality in Nature: Structure– Function Relationships in Biological Materials. Biomimetics 8, e284 (2023).

3. J. Michael, J. McFarland, Another look at the core concepts of physiology: revisions and resources. Adv Physiol Educ. 44(4):752–762 (2020) doi: 10.1152/advan.00114.2020

4. I. Koubiyr, M. Deloire, B. Brochet, P. Besson, J. Charré-Morin, A. Saubusse, T. Tourdias, A. Ruet, Structural constraints of functional connectivity drive cognitive impairment in the early stages of multiple sclerosis. Mult Scler. 27(4):559–567 (2021). doi: 10.1177/1352458520971807.

5. Z. Wu, S. Feng, K. Li, L. Dong, L. Zhang, Y. Ning, D. Yin, Altered Structure-Function Coupling Associated with Attention Decline in Shift Work Disorder. Nat Sci Sleep. 21;17:989–1001 (2025). doi: 10.2147/NSS.S503303.

6. S.S. Sivan, E. Wachtel, P. Roughley Structure, function, aging and turnover of aggrecan in the intervertebral disc, Biochim Biophys Acta. 1840(10):3181–9 (2014). doi: 10.1016/j.bbagen.2014.07.013.

7. J.B. Strait, E.G. Lakatta, Aging-associated cardiovascular changes and their relationship to heart failure. Heart Fail Clin. 8(1):143–64 (2012) doi: 10.1016/j.hfc.2011.08.011. PMID: 22108734

8. H. Schulte, C. Mühlfeld, C. Brandenberger Age-Related Structural and Functional Changes in the Mouse Lung. Front Physiol. 4;10:1466 (2019). doi: 10.3389/fphys.2019.01466. eCollection 2019. PMID: 31866873

9. R. Berlin, R. Gruen, J.J. Best, Systems Medicine-Complexity Within, Simplicity Without. Healthc Inform Res. 1(1):119–137 (2017). doi: 10.1007/s41666-017-0002-9. Epub 2017 May 10. PMID: 28713872

10. S.D. Castle, C.S. Grierson, T.E. Gorochowski, Towards an engineering theory of evolution. Nat Commun. 2021 Jun 7;12(1):3326. doi: 10.1038/s41467-021-23573-3. PMID: 34099656

11. P. Liberali, A.F. Schier, The evolution of developmental biology through conceptual and technological revolutions, Cell, 187(14):3461–3495

12. J. Kim, Making sense of emergence. Philos Stud 1999 95:3–36

13. M.H.V. Van Regenmortel, D. Hull Promises and Limits of Reductionism in the Biomedical Sciences. Wiley, Chichester, UK (2002)

14. M.H.V. Van Regenmortel Reductionism and complexity in molecular biology. EMBO Rep 5(11):1016–1020 (2004)

15. A.J. Ma’ayan, Complex systems biology. R Soc Interface. 14(134):20170391 (2017) doi: 10.1098/rsif.2017.0391

16. R.V. Solé, B.C. Goodwin, Signs of Life: How Complexity Pervades Biology. New York: Basic Books (2000)

17. A. Dambricourt Malassé, Self-Organization as a New Paradigm in Evolutionary Biology, Springer Cham series Evolutionary Biology – New Perspectives on Its Development, 10.1007/978-3-031-04783-1

18. M. Adler, A.R. Chavan, R. Medzhitov, Tissue Biology: In Search of a New Paradigm. Annu Rev Cell Dev Biol. 39:67–89 (2023). doi: 10.1146/annurev-cellbio-120420-113830

19. A.A. Cohen, L. Ferrucci, T. Fülöp, D. Gravel, N. Hao, A. Kriete, M.E. Levine, L.A. Lipsitz,M.G.M. Olde Rikkert, A. Rutenberg, N. Stroustrup, R. Varadhan, A complex systems approach to aging biology. Nat Aging. 2(7):580–591 (2022). doi: 10.1038/s43587-022-00252-6. Epub 2022 Jul 20. PMID: 37117782

20. N. Claussen, F. Brauns, S. J. Streichan; Searching for physical principles of morphogenesis. Development 152 (21):dev204894 (2025). doi: 10.1242/dev.20489

21. R. López-Ruiz, Mathematical Biology: Modeling, Analysis, and Simulations. Mathematics 10, e3892 (2022).

22. B. Breckling, Individual-based modelling: potentials and limitations. ScientificWorldJournal. 19;2:1044–62 (2002) doi: 10.1100/tsw.2002.179

23. D. Drasdo, On Selected Individual-based Approaches to the Dynamics in Multicellular Systems, in Polymer and Cell Dynamics: Multiscale Modeling and Numerical Simulations, W. Alt, M. Chaplain, M. Griebel, J. Lenz, Eds. (Birkhäuser, 2003), pp. 169–203.

24. J. Galle, M. Ho mann, G. Aust, From single cells to tissue architecture : a bottom-up approach to modelling the spatio-temporal organization of complex multicellular systems. J Math Biol 58(1-2):261–83 (2009)

25. M. Wang, M. Garbey, S. A. Berceli, R. Tran-Son-Tay, Rule-Based Simulation of Multi-Cellular Biological Systems – A Review of Modeling Techniques. Cellular and Molecular Bioengineering 2, 285–294 (2009)

26. J.D. Humphrey, E.R. Dufresne, M.A. Schwartz, Mechanotransduction and extracellular matrix homeostasis. Nat. Rev. Mol. Cell Biol. 15, 802–812 (2014).

27. Z. Sun, S.S. Guo, R. Fässler Integrin-mediated mechanotransduction. J Cell Biol. 215(4):445–456. (2016) doi: 10.1083/jcb.201609037

28. A. Totaro, T. Panciera, S. Piccolo, YAP/TAZ upstream signals and downstream responses. Nat Cell Biol. 20(8):888–899. (2018) doi: 10.1038/s41556-018-0142-z

29. A. Pocaterra, P. Romani, S. Dupont YAP/TAZ functions and their regulation at a glance. J Cell Sci. 133(2):jcs230425. doi: 10.1242/jcs.230425 (2020)

30. A. Saraswathibhatla, D. Indana, O. Chaudhuri, Cell-extracellular matrix mechanotransduction in 3D. Nat Rev Mol Cell Biol. 24(7):495–516. (2023) doi: 10.1038/s41580-023-00583-1

31. P. Romani, L. Valcarcel-Jimenez, C. Frezza, S. Dupont, Crosstalk between mechanotransduction and metabolism. Nat Rev Mol Cell Biol 22, 22–38 (2021). 10.1038/s41580-020-00306-w

32. L. Le Goff, T. Lecuit, Mechanical Forces and Growth in Animal Tissues, Cold Spring Harb Perspect Biol. 8(3):a019232 (2015) doi: 10.1101/cshperspect.a019232

33. R.J. Nims, L. Pferdehirt, F. Guilak, Mechanogenetics: harnessing mechanobiology for cellular engineering. Curr Opin Biotechnol 73:374–379 (2022) doi: 10.1016/j.copbio.2021.09.011.

34. Z. Qin, N.M. Pugno, M.J. Buehler, Mechanics of fragmentation of crocodile skin and other thin films. Sci Rep. 4:4966. (2014) doi: 10.1038/srep04966

35. A.J. Richard, U. White, C.M. Elks, J.M. Stephens, Adipose Tissue: Physiology to Metabolic Dysfunction. (2020) In: Feingold KR, Ahmed SF, Anawalt B, Blackman MR, Boyce A, Chrousos G, Corpas E, de Herder WW, Dhatariya K, Dungan K, Hofland J, Kalra S, Kaltsas G, Kapoor N, Koch C, Kopp P, Korbonits M, Kovacs CS, Kuohung W, Laferrère B, Levy M, McGee EA, McLachlan R, Muzumdar R, Purnell J, Rey R, Sahay R, Shah AS, Singer F, Sperling MA, Stratakis CA, Trence DL, Wilson DP, editors. Endotext [Internet]. South Dartmouth (MA): MDText.com, Inc.; 2000–. PMID: 32255578

36. N. Alkhouli, J. Mansfield, E. Green, J. Bell, B. Knight, N. Liversedge, J. C. Tham, R. Welbourn, A. C. Shore, K. Kos, and C.P. Winlove, The mechanical properties of human adipose tissues and their relationships to the structure and composition of the extracellular matrix, American Journal of Physiology-Endocrinology and Metabolism 305:12, E1427–E1435 (2013)

37. J. Dichamp, C. Barreau, C. Guissard, A. Carrière, Y. Martinez, X. Descombes, L. Pénicaud, J. Rouquette, L. Casteilla, F. Plouraboué, A. Lorsignol 3D analysis of the whole subcutaneous adipose tissue reveals a complex spatial network of interconnected lobules with heterogeneous browning ability. Sci Rep. 9(1):6684 (2019). doi: 10.1038/s41598-019-43130-9. PMID: 31040317

38. F. Wassermann, “The development of adipose tissue” in Comprehensive Physiology, R. Terjung, Ed. (John Wiley & Sons, 2011), pp. 87–100

39. M. Crisan, S. Yap, L. Casteilla, C.W. Chen, M. Corselli, T.S. Park, G. Andriolo, B. Sun, B. Zheng, L. Zhang, C. Norotte, P.N. Teng, J. Traas, R. Schugar, B.M. Deasy, S. Badylak, H.J. Buhring, J.P. Giacobino, L. Lazzari, J. Huard, B. Péault A perivascular origin for mesenchymal stem cells in multiple human organs, Cell Stem Cell, 3, pp. 301–313 (2008)

40. W. Tang, D. Zeve, J.M. Suh, D. Bosnakovski, M. Kyba, R.E. Hammer, M.D. Tallquist, J.M. Graff White fat progenitor cells reside in the adipose vasculature Science, 322 pp. 583–586 (2008)

41. G. Ailhaud Cross talk between adipocytes and their precursors: relationships with adipose tissue development and blood pressure Ann. N. Y. Acad. Sci., 892 pp. 127–133 (1999)

42. M. Hemmingsen, et al. The role of paracrine and autocrine signaling in the early phase of adipogenic differentiation of adipose-derived stem cells PLoS ONE, 8 (5) p. je63638 (2013)

43. P. Chassonnery, J. Paupert, A. Lorsignol, C. Sévérac, M. Ousset, P. Degond, L. Casteilla, D. Peurichard, Fiber crosslinking drives the emergence of order in a three-dimensional dynamical network model. Royal Society Open Science 11, 231456 (2024).

44. S. C. Cowin, Tissue Growth and Remodeling. Annual Review of Biomedical Engineering 6, 77– 107 (2004).

45. M Abedin, N King, Diverse evolutionary paths to cell adhesion. Trends Cell Biol. 12:734–42 (2010). doi: 10.1016/j.tcb.2010.08.002.

46. JM Tarbell, LM Cancel. The glycocalyx and its significance in human medicine. J Intern Med. 280(1):97–113 (2016). doi: 10.1111/joim.12465

47. O. Campàs, I. Noordstra, A.S. Yap, Adherens junctions as molecular regulators of emergent tissue mechanics. Nat Rev Mol Cell Biol. 25(4):252–269 (2024) doi: 10.1038/s41580-023-00688-7

48. R.J. Johnston, C. Desplan, Stochastic mechanisms of cell fate specification that yield random or robust outcomes. Annu Rev Cell Dev Biol. 26:689–719 (2010) doi: 10.1146/annurev-cellbio-100109-104113

49. SJ Gould, RC Lewontin The spandrels of San Marco and the Panglossian paradigm: a critique of the adaptationist programme. Proc R Soc Lond B Biol Sci. 1979 Sep 21;205(1161):581–98. doi: 10.1098/rspb.1979.0086.

50. S.A. Newman, G. Forgacs, G.B. Muller, Before programs: the physical origination of multicellular forms. Int J Dev Biol. 50(2-3):289–99 (2006) doi: 10.1387/ijdb.052049sn.

51. M.N. Shahbazi, E.D. Siggia, M. Zernicka-Goetz, Self-organization of stem cells into embryos: A window on early mammalian development. Science. 364(6444):948–951 (2019) doi: 10.1126/science.aax0164. PMID: 31171690; PMCID: PMC8300856.

52. M. Zhu, M. Zernicka-Goetz, Principles of Self-Organization of the Mammalian Embryo. Cell 183(6):1467–1478 (2020) doi: 10.1016/j.cell.2020.11.003. PMID: 33306953; PMCID: PMC8212876.

53. D.E. Ingber, The origin of cellular life, Bioessays 22(12):1160–70 (2000) doi: 10.1002/1521-1878(200012)22:12<1160::AID-BIES14>3.0.CO;2-5.

54. S.A. Newman, Physico-genetic determinants in the evolution of development. Science 338(6104):217–9 (2012) doi: 10.1126/science.1222003. PMID: 23066074

55. J. Schindelin, I. Arganda-Carreras, E. Frise, V. Kaynig, M. Longair, T. Pietzsch, S. Preibisch, C. Rueden, S. Saalfeld, B. Schmid, J. Tinevez, D. J. White, V. Hartenstein, K. Eliceiri, P. Tomancak, A. Cardona, Fiji: an open-source platform for biological-image analysis. Nature Methods 9, 676–682 (2012).

56. S. Preibisch, S. Saalfeld, P. Tomancak, Globally optimal stitching of tiled 3D microscopic image acquisitions. Bioinformatics 25, 1463–1465 (2009).

57. C. Stringer, T. Wang, M. Michaelos, M. Pachitariu, Cellpose: a generalist algorithm for cellular segmentation. Nature Methods 18, 100–106 (2021).

58. M. Pachitariu, C. Stringer, Cellpose 2.0: how to train your own model. Nature Methods 19, 1634–1641 (2022).

59. M Linkert et al. Metadata matters: access to image data in the real world. Journal of Cell Biology 189(5), 777–782 (2010)

60. M Berth et al., The state of the art in the analysis of two-dimensional gel electrophoresis images. Appl. Microbiol Biotechnol 76, 1223–1243 (2007)

61. F. Meyer, Topographic distance and watershed lines. Signal Processing 38, 113–125 (1994).

62. J. B. T. M. Roerdink, A. Meijster, The Watershed Transform: Definitions, Algorithms and Parallelization Strategies. Fundamenta Informaticae 41, 187–228 (2000).

63. J. Ahrens, B. Geveci, C. Law, “ParaView: An End-User Tool for Large Data Visualization” in Visualization Handbook, C. D. Hansen, C. R. Johnson, Eds. (Elsevier, 2005), pp. 717–731.

