## Supplementary material for "Emergence and robustness of 3D complex tissue architectures from simple physical forces": SI text

**This PDF file includes:**

Figures S1 to S5
Table S1

**Fig. S1.**
**Segmentation of *in-vivo* samples. Related to  1 of the main text**

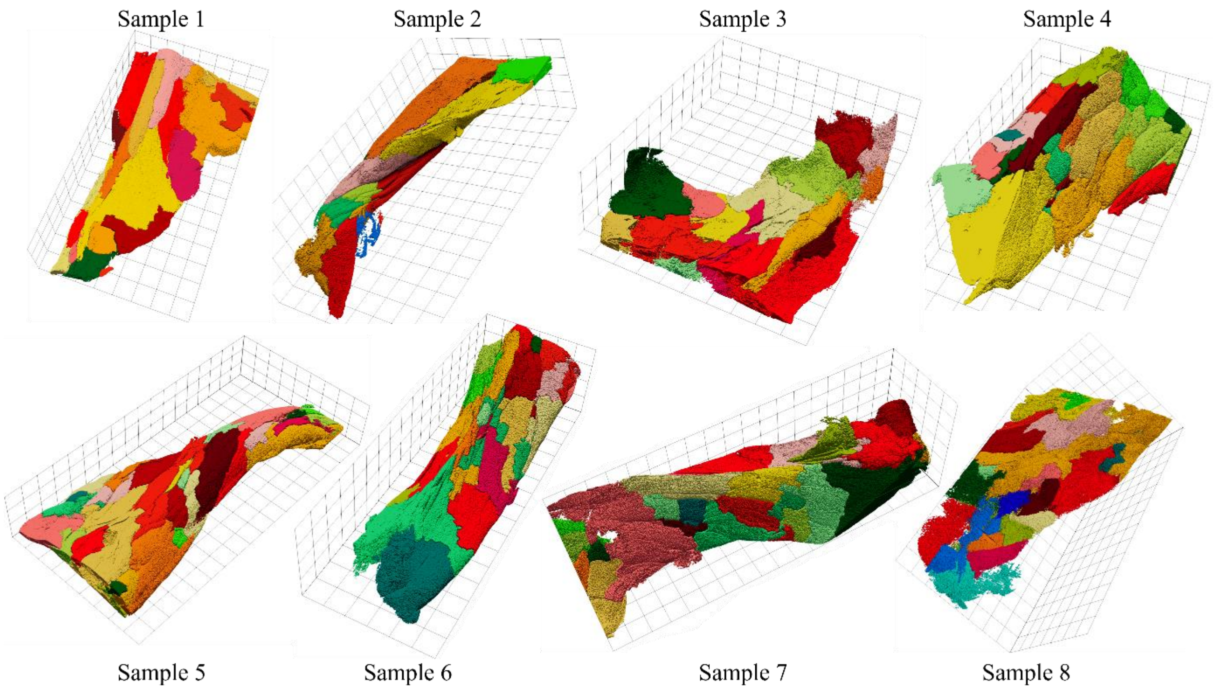

**Fig. S2.**

**a.** Cells (left) and fibers (right). **b.** Biological phenomena: cell insemination and growth (left) and fiber linking/unlinking (right). **c.** Mechanical interactions: from left to right: cell-cell, fiber-cell, fiber-fiber repulsions, fiber alignment. **d.** Cell insemination probability as function of parameter $b_{ins}$.

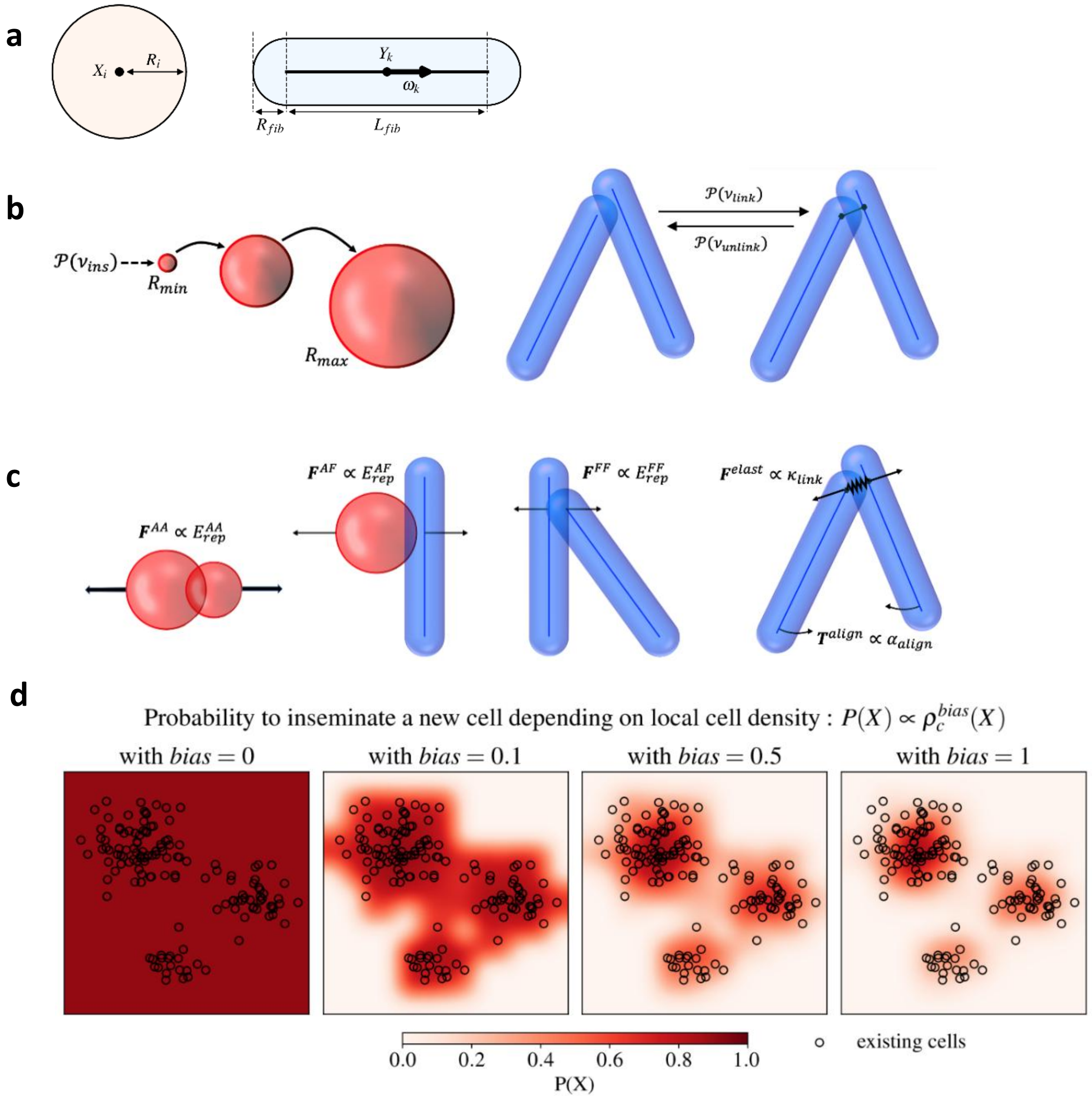

**Fig. S3.**

**a.** Distribution of the *in silico* cell clusters (total 266 420 clusters from 12600 simulations) according to the morphological quantifiers $E_{lob}$ (elongation, in ordinate) and $S_{lob}$ (solidity, in abscissa). Red asterisks are related to the three clusters highlighted in Fig. 2.a. **b.** Values of the *in silico / in vivo* distance $\Delta Q_{sample}$ in the 4D plane $(\phi_{fib}, \phi_{ad}, \chi_{link}, \nu_{link})$ (from black ($\Delta Q_{sample} \ll 1$) to white ($\Delta Q_{sample} > 20$). For a pair value of $\phi_{ad}$ (columns) and $\phi_{fib}$(lines), each heatmap gives the values of $\Delta Q_{sample}$ as function of the ECM crosslinking rate $\nu_{link}$ (in ordinate) and ECM connectivity $\chi_{link}$ (in abscissa), averaged over 10 simulations. **c.** Scatter plot of the SHAP value for $\nu_{link}$ with its main interaction feature $\chi_{link}$, with a sigmoid function fitted by non-linear least squares. Large values of the interaction parameter are in red, small values in blue. **d.** Scatter plot of the SHAP value for $\phi_{ad}$ colored according to its main interaction feature $\chi_{link}$. Large values of the interaction parameter are in red, small values in blue.

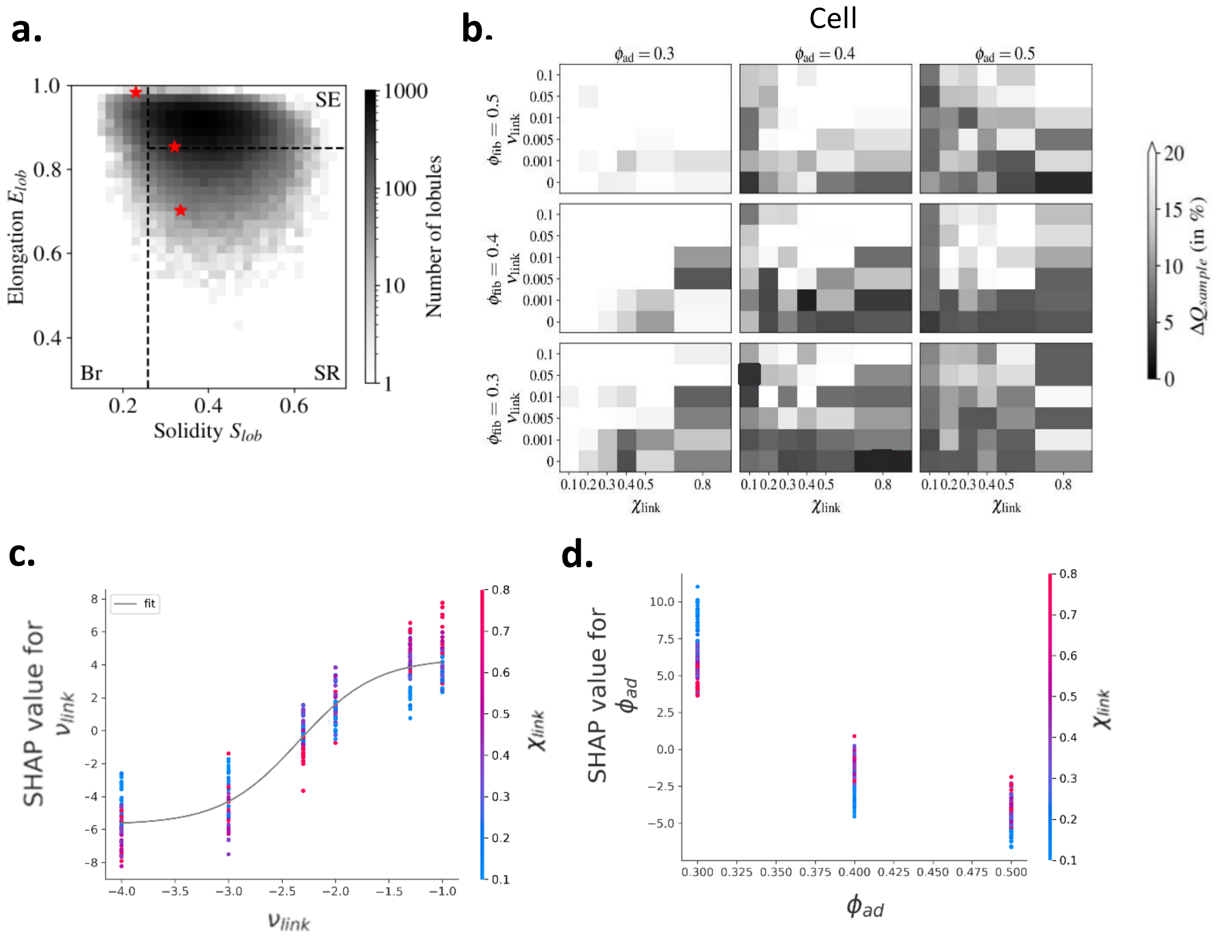

**Fig. S4**

**a.** Simulations at equilibrium for $\Phi_{fib} = 0.4$, lowly crosslinked ECM $\chi_{link} = 0.1$. **top line**: $\Phi_{ad} = 0.4$, **bottom line**: $\Phi_{ad} = 0.3$. **left column**: low crosslinking rate $\nu_{link} = 10^{-3}$. **right column**: fast crosslinking rate $\nu_{link} = 0.1$. Cells are colored according to their cluster category (red/yellow for SE, green for SR, blue for Br). Fibers are colored according to their local alignment (from blue (low) to red (large)). The SE/SR/Br proportions for the 10 simulations of the selected parameter sets and their weighted mean are indicated in the bottom bar plots. **b. top**: mean (weighted) proportions of SE (red), SR (green) and Br (blue) clusters as function of $\nu_{link}$ (log10 scale) for $\Phi_{ad} = 0.4$ (plain curves) and $\Phi_{ad} = 0.3$ (dotted curves), and for *in vivo* data (shaded error bars). Plain error bars indicate weighted standard deviation between simulations (10 simulations per parameter values). **bottom**: Fiber mean alignment as function of $\nu_{link}$ (log10 scale).

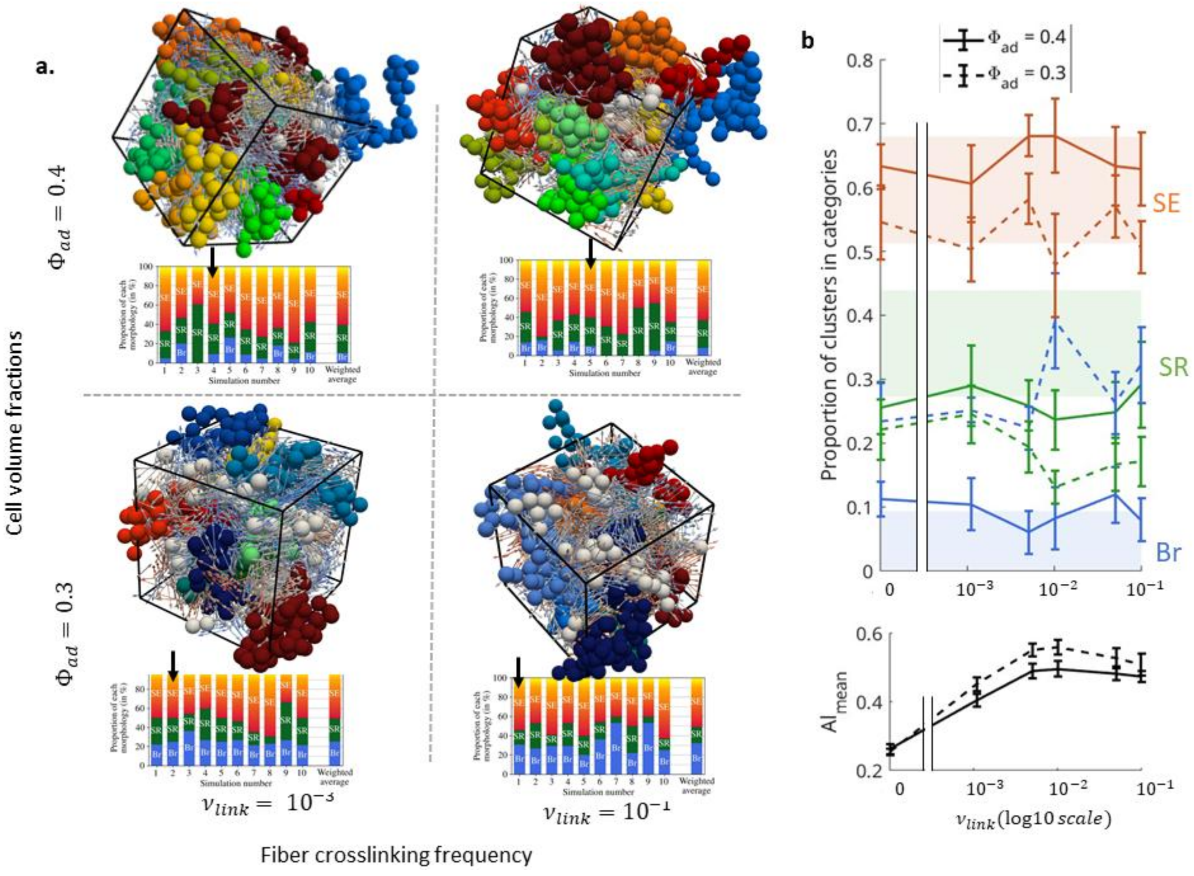

**Fig. S5**

**a.** Absolute mean plot highlighting interaction effects (off-diagonal) of parameters in the prediction of $\Delta Q_{sample}$. Colors represent large (dark red) to small (dark blue) effects. **b.** Scatter plot of the SHAP value for $\phi_{ad}$colored according to its main interaction feature $b_{ins}$ (large values in red, small values in blue). **c.** Boxplots of the number of cell clusters as functions of of the bias parameter $b_{ins}$ (abscissa), all other parameters combined. **From left to right**: total number of cell clusters, number of SE clusters, number of SR clusters, number of Br clusters.

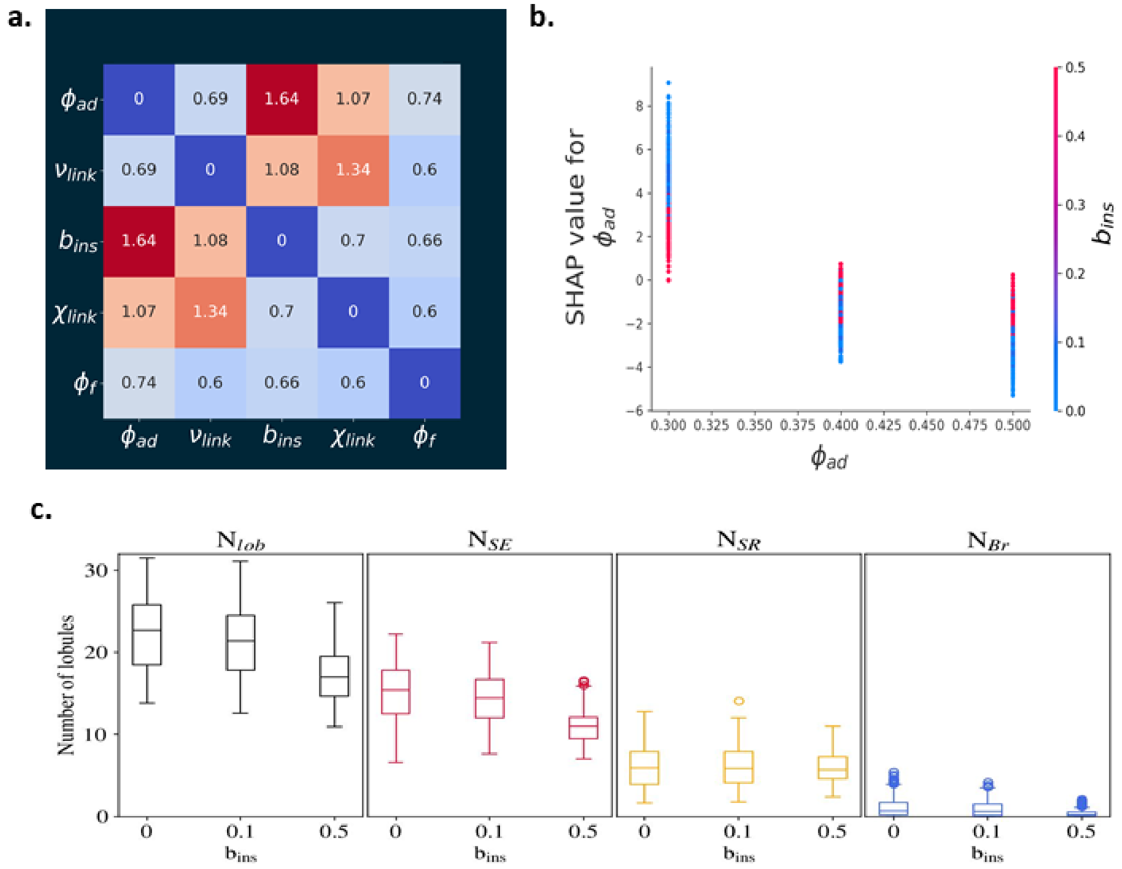

**Table S1.**
**Summary of the model parameters**

| Name | Value | Units | Description |
|---|---|---|---|
| **Agents** | | | |
| $\phi_{ad}$ | [0.3, 0.5] | N/A | Maximum cell volume fraction |
| $R_{ad}^{min}$ | 0.1 | L | Minimum cell radius (at insemination) |
| $R_{ad}^{max}$ | 1.5 | L | Maximal cell radius |
| $\phi_{fib}$ | [0.3, 0.5] | N/A | Fiber volume fraction |
| $L_{fib}$ | 6 | L | Fiber unit's length |
| $R_{fib}$ | 0.5 | L | Fiber unit's radius |
| **Mechanical interactions** | | | |
| $\alpha_{rep}^{AA}$ | 5 | $M\ L^{-1}\ T^{-2}$ | Cell-cell repulsion intensity |
| $\alpha_{rep}^{AF}$ | 8.9 | $M\ L^{-1}\ T^{-2}$ | Cell-fiber repulsion intensity |
| $\alpha_{rep}^{FF}$ | 12.5 | $M\ L^{-1}\ T^{-2}$ | Fiber-fiber repulsion intensity |
| $\alpha_{rest}$ | 5 | $M\ T^{-2}$ | Elastic crosslink restoring force intensity |
| $\alpha_{align}$ | 2 | $ML^2\ T^{-2}$ | Elastic crosslink alignment intensity |
| $\mu_{ad}$ | 1 | $ML^{-1}T^{-1}$ | Cell dynamic viscosity |
| $\mu_{fib}$ | 1 | $ML^{-1}T^{-1}$ | Fiber units dynamic viscosity |
| **Cell processes** | | | |
| $\nu_{ins}$ | 1 | $T^{-1}$ | Cell insemination frequency |
| $b_{ins}$ | [0, 1] | N/A | Biased insemination parameter |
| $K_{growth}$ | 0.05 | $L^3T^{-1}$ | Cell volume growth per unit time |
| **Fiber crosslink processes** | | | |
| $\nu_{link}$ | [0.001, 0.1] | $T^{-1}$ | Fiber crosslinking frequency |
| $\chi_{link}$ | [0.1, 0.8] | N/A | Linked fiber fraction at equilibrium |
| $d_{link}^{max}$ | 1.2 | L | Perception distance for crosslink creation |
| $d_{link}^{eq}$ | 1 | L | Crosslink equilibrium length |
| **Numerical parameters** | | | |
| $L_x = L_y = L_z$ | 30 | L | Side length of the cubic domain |
| $T_{final}$ | 2000 | T | Total simulation time |
